# Cholesterol binding to the transmembrane region of a group 2 HA of Influenza virus is essential for virus replication affecting both virus assembly and HA’s fusion activity

**DOI:** 10.1101/598953

**Authors:** Bodan Hu, Chris Tina Höfer, Christoph Thiele, Michael Veit

## Abstract

Hemagglutinin (HA) of Influenza virus is incorporated into cholesterol enriched, nanodomains of the plasma membrane. Phylogenetic group 2 HAs contain the conserved cholesterol consensus motif (CCM) YKLW in the transmembrane region. We previously reported that mutations in the CCM retarded intracellular transport of HA and decreased its nanodomain association. Here we analyzed whether cholesterol interacts with the CCM. Incorporation of photocholesterol into HA was significantly reduced if the whole CCM is replaced by alanine, both using immunoprecipitated HA and when HA is embedded in the membrane. Next, we used reverse genetics to investigate the significance of the CCM for virus replication. No virus was rescued if the whole motif is exchanged (YKLW4A); single (LA) or double (YK2A and LW2A) mutated virus showed decreased titers and a comparative fitness disadvantage. In polarized cells transport of HA mutants to the apical membrane was not disturbed. Reduced amounts of HA and cholesterol were incorporated into the viral membrane. Mutant viruses exhibit a decrease in hemolysis, which is only partially corrected if the membrane is replenished with cholesterol. More specifically, viruses have a defect in hemifusion as demonstrated by fluorescence dequenching. Cells expressing HA-YKLW4A fuse with erythrocytes, but the number of events are reduced. Even after acidification unfused erythrocytes remain cell-bound, a phenomenon not observed with wildtype HA. We conclude that cholesterol-binding to a group 2 HA is essential for virus replication. It has pleiotropic effects on virus assembly and membrane fusion, mainly on lipid mixing and possibly a preceding step.

**IMPORTANCE:** The glycoprotein hemagglutinin (HA) is a major pathogenicity factor of Influenza viruses. Whereas the structure and function of HA’s ectodomain is known in great detail, similar data for the membrane-anchoring part of the protein are missing. Here we demonstrate that the transmembrane region of a group 2 HA interacts with cholesterol, the major lipid of the plasma membrane and the defining element of the viral budding site nanodomains of the plama membrane. The cholesterol binding motif is essential for virus replication. Its partial removal affects various steps of the viral life cycle, such as assembly of new virus particles and their subsequent cell entry via membrane fusion. A cholesterol-binding pocket in group 2 HAs might be a promising target for a small lipophilic drug that inactivates the virus.

## INTRODUCTION

Hemagglutinin (HA) of Influenza virus is a typical type I transmembrane glycoprotein with an N-terminal signal peptide, a large ectodomain, a single transmembrane region and a short cytoplasmic tail (1). HA assembles into homotrimers in the ER and is transported via the secretory pathway to the plasma membrane, in polarized cells to the apical membrane, where virus assembly and budding take place (2). It was proposed that Influenza virus assembles in and buds through small dynamic, cholesterol- and sphingolipid-enriched nanodomains of the plasma membrane which could coalesce to larger, more stable platforms (3, 4). Indeed, it was demonstrated by quantitative mass spectrometry that these lipids are enriched in the viral membrane relative to the entire apical membrane of their host cell (5). HA organizes the viral assembly site, since it is not randomly distributed in the plasma membrane of transfected cells, but is present in (partly cholesterol-sensitive) clusters of various sizes as demonstrated by quantitative immunoelectron microscopy (6–8) and FPALM (fluorescence-photoactivation-localization microscopy) (9). The other integral membrane proteins of the virus, the neuraminidase (NA) and the ion channel M2 contain their own signals for targeting to the viral assembly site (10), where they are supposed to recruit the internal components of viral particles, the matrix protein M1 and the eight ribonucleoparticles (containing the viral genome segments complexed to the nucleoprotein (NP) and the three polymerase proteins PA, PB1 and PB2) into budding virions (11, 12).

HA plays also a pivotal role during virus entry. It is responsible for receptor recognition: A binding pocket in the globular head domain of the molecule recognizes sialic acid moieties in glycoproteins and glycolipids on the host cell surface. After clathrin-mediated endocytosis of the virus acidification of the endosome triggers an irreversible conformational change in HA (1). In order to perform the conformational change the inactive precursor HA0 must first be processed into two disulfide-linked subunits, the membrane-embedded HA2 and the globular HA1 subunit, by a protease provided by the host organism (13). The elucidation of the crystal structures of the ectodomain of HA at neutral and a part of the ectodomain at mildly acidic pH has led to a model of how conformational changes of HA execute membrane fusion. The hydrophobic fusion peptide at the N-terminus of HA2, which is buried inside the trimeric structure at neutral pH, becomes exposed on the distal end of the molecule after acidification and interacts with the cellular membrane. A second conformational change then bends the HA-molecule thereby drawing the fusion peptide towards the transmembrane region which leads to close apposition of both lipid bilayers (1). By mechanisms that are not well understood at present lipid exchange initially only occurs between the outer leaflets of the viral and the cellular membrane (hemifusion). Finally, a fusion pore opens, flickers and dilates thereby allowing entry of the viral genome into the target cell (14). Analysing the fusion kinetics of individual virus particles indicated that exposure of the fusion peptide is the rate-limiting step of the reaction. Full fusion then requires the cooperative action of three to four neighbouring HA trimers that pull together on the cellular membrane to execute fusion (15).

The C-terminal membrane anchoring fragment of HA, for which no structure has been elucidated, also contributes to membrane fusion. Fusion pore formation requires the presence of a transmembrane region with a minimal length of 17 amino acids (16); HA anchored to the outer leaflet of the membrane by a glycolipid instead of the TMR may cause hemifusion after acidification, but is not able to catalyze full fusion (17). The cytoplasmic tail plays a role during fusion pore formation; it requires S-acylation at conserved cysteine residues at least in some HA subtypes (18), whereas other (artificial) modifications of the tail negatively affect fusion (19, 20).

The lipids in the membrane destined to fuse are also not passive bystanders of the reaction. Since fusion requires strong bending of the bilayer, certain lipid species with an intrinsic curvature, i.e. those having a small head group and a large tail (or *vice versa*) positively or negatively affect certain stages of the reaction (21). One abundant lipid species with a negative intrinsic curvature is cholesterol, since it consists of a small hydrophilic head group, a large and rigid steroid ring structure and a flexible hydrocarbon tail and thus it may stabilize highly curved fusion intermediates. Indeed, cholesterol addition or removal from the HA-containing membrane positively or negatively affects the extent of fusion (22, 23). Cholesterol acts at two stages in membrane fusion: at an early, lipidic stage prior to fusion pore opening and a later stage during fusion pore expansion (22). However, the mechanism of the effect of cholesterol on fusion is not understood and is probably more complex than stabilizing highly curved lipid intermediates. Cholesterol prevents unphysiological (“leaky”) fusion reactions (24), affects membrane ordering (and hence the lateral mobility of HA), the spatial distance of HA in virus particles (25) and finally allows liquid phase separation that concentrates HA for efficient fusion (26).

Hemagglutinin has two signals for targeting to cholesterol-enriched nanodomains. On one hand, three conserved S-acylated cysteines located at the cytoplasmic end of the TMR and in the cytoplasmic domain, respectively (27–29), on the other hand, hydrophobic amino acids in the TMR facing the outer leaflet of the plasma membrane, especially the conserved amino acids VIL at the beginning of the TMR. The latter were identified by alanine scanning mutagenesis throughout the whole TMR of HA to identify residues that confer incorporation of HA into detergent-resistant-membranes (DRMs), the biochemical correlate of membrane nanodomains (26, 30). Mutation of these amino acids at the beginning of the TMR also reduced fluorescence resonance energy transfer (FRET) of HA with a double-acylated raft-marker (27, 29, 31).

We reported recently that leucine of VIL might be part of a cholesterol consensus motif (CCM) that is known to bind cholesterol to 7-transmembrane-receptors (7TMR). By crystallography the cholesterol-interacting amino acids in the human ß-adrenergic receptor were identified and by sequence comparison with other 7TMR the cholesterol consensus motif (CCM) was defined (32). In the CCM the amino acids interacting with cholesterol are not a linear sequence motif, but distributed between two transmembrane helices of these polytopic membrane proteins. One helix contains the sequence motif W/Y-I/V/L-K/R, whereby all residues must face the same side of the helix. In addition, another aromatic amino acid, either phenylalanine (F) or tyrosine (Y) is needed on a second helix to bind cholesterol from the other side. Since cholesterol is present in both leaflets of a bilayer (33), the CCM can be orientated in two ways; the charged amino acid might face either the extracellular space or the cytosolic compartment (34–36).

HAs of the phylogenetic group 2 contain a strictly conserved YKLW motif which conforms to the CCM defined for 7TMR (F/Y-R/K-I/V/L-Y/W). Mutations in the CCM drastically retard Golgi-localized processing of HA, such as acquisition of Endo-H resistant carbohydrates in the medial-Golgi and proteolytic cleavage in the TGN. All mutants analysed by FRET also showed reduced association with nanodomains at the plasma membrane (37).

Here we analysed whether the CCM indeed interacts with cholesterol and if mutations in the CCM affect virus replication, virus assembly and membrane fusion.

## MATERIAL AND METHODS

### Cell culture and virus experiments

Madin Darby canine kidney (MDCK II), Chinese hamster ovary (CHO) and human embryonic kidney 293T cells were grown in DMEM (Dulbecco’s modification of Eagle’s medium, PAN, Aidenbach, Germany) supplemented with 10% FCS (fetal calf serum, Perbio, Bonn, Germany) and penicillin/streptomycin (100 units/ml and 100 μg/ml, respectively) at 37 °C and 5% CO2. To generate polarized cells, 5×10 MDCK II cells were seeded into 24 mm transwells containing a polyester membrane with pores having a diameter of 0.4 µm (Corning) using 1.5 ml growth medium in the upper and 2.6 ml in the lower chamber. Medium was exchanged every day and cells were cultured for 4 days.

Mutant 1 of the highly pathogenic strain A/FPV/Rostock/1934 (H7N1), termed FPV*, that contains the sequence PSKGR instead of PSKKRKKR at the C-terminus of HA1 (38) was used to create recombinant virus. FPV* shows low pathogenicity in chicken and requires trypsin for growth in cell culture and is thus suitable for working in a BSL2 lab. The full-length sequence (excluding the fluorophore) of HA mutants LA, YK2A, LW2A and YKLW4A were cloned from plasmid pECerulean (37) to pHH21 with In-Fusion® HD Cloning Kit (Takara Bio, Japan). Recombinant Influenza viruses were produced with the twelve plasmids system (38) by transfection (0.5 µg of each plasmid) of 293T cells in 35 mm dishes with TurboFect reagent. 4-6 h later, the medium was changed to infection medium (DMEM, 0.1% FCS, 0.2% BSA, 1 µg/ml TPCK-Trypsin). 48 h post transfection, the supernatant was harvested and centrifuged at 2000 g for 5 min to clear from cell debris and further amplified in MDCK II cells to generate a virus stock.

HA tests were performed in 96 well U-bottom microwell plates. 50 µl of a two-fold serial dilution of virus sample in PBS was incubated with 50 µl 1% chicken red blood cells for 30 minutes at room temperature.

For TCID50 tests virus samples were three-fold serially diluted in infection medium. 100 µl diluted virus samples (8 replicates) were added to confluent MDCK II cells in 96-well plates after washing the cells once with DPBS+ (Dulbecco’s phosphate-buffered saline with Calcium and Magnesium, PAN biotech, Germany). 2 days post infection, cells were washed once with PBS, fixed with 3% formaldehyde in PBS for 5 min and stained with 0.1% crystal violet. The TCID50 titer was calculated by Reed & Muench Method.

To generate a growth curve, 90% confluent MDCK II cells in 6-well plates were infected with FPV*-wt or mutants with a moi of 0.0005 (based on TCID50 titer). 1 h post infection, medium was replaced by infection medium, aliquots of culture supernatant were harvested at 12 h, 24 h, 36 h and 48 h post infection, cleared from cell debris (2000 g, 5 min) and titrated by TCID50 and HA-assay.

For competitive growth experiments FPV* mutant was mixed with wild type at a ratio of 5 to 1 and MDCK II cells were infected with total moi of 0.0005. vRNA was extracted from the virus mixture or at 24 h and 48 h post infection from the cell culture supernatant with RTP® DNA/RNA Virus Mini Kit (Stratec, Germany). OneStep RT-PCR Kit (Qiagen) was used for reverse transcription of HA fragment with specific primers (Forward: TGAAAATGGTTGGGAAGGTCTGG, Reverse: CGCATGTTTCCGTTCTTCACAC), which were then sent for sequencing.

For growth experiments in polarized MDCK II cells, they were infected with viruses through the upper chamber at an m.o.i. of 0.001. After binding for 1 h, the culture medium in the upper chamber was changed to infection medium. The culture medium from upper and lower chamber was harvested separately at 8 h, 24 h, 34 h and 48 h post infection and virus titer was determined by HA-assay.

### Photocholesterol crosslinking of HA

Two different experiments, labeling of HA-expressing cells or immunoprecipitated HA, were performed to investigate whether HA-wt and HA with mutations in the CCM interact with click-photocholesterol (6,6’-Azi-25-ethinyl -27-norcholestan-3ß-ol, see supplementary file 1 for synthesis of the compound). For the first experiment, CHO cells in 6-well plates were transfected with HA-wt, HA-LA, HA-YK2A, HA-LW2A or HA-YKLW4A cloned into the vector pCAGGS. 4-6 h post transfection, 5 µl photocholesterol (from 5 mg/ml stock in ethanol, final concentration is 50 µM) was added to 1ml of medium without serum and cells were incubated overnight. 24 hours after transfection cells were put on ice and exposed to UV light (wavelength 320–365 nm, power 8W, 3.5 A, 60V) for 10 minutes to activate the diazirine group. Cells were then lysed in 500 µl 1% NP40 in IP buffer (500 mM Tris-HCl, 20 mM EDTA, 30 mM sodium pyrophosphate decahydrate, 10 mM sodium fluoride, 1 mM sodium orthovanadate, 2 mM benzamidine, 1 mM PMSF, 1 mM NEM and protease inhibitor cocktail (Sigma)). 450 µl (=90%) of the cell lysate was incubated with anti-HA2 antiserum (1:1000) at 4°C with agitation overnight. 50 µl of protein-A-sepharose was added to the mixture and incubated at 4°C for another 4h prior to pelleting and washing with IP-buffer (2x) and with PBS.

For cholesterol crosslinking of purified protein, HA was first immunoprecipitated from transfected CHO cells, 0.5 µl photocholesterol was added to immunoprecipitated HA (in 100 µl PBS) and the mixture was illuminated with UV light at 4°C for 10 minutes.

HA-photocholesterol complexes were then clicked to Pico-azido picolyl sulfo cy3 by using the CuAAC Biomolecule Reaction Buffer Kit (Jena Bioscience). Samples were subjected to SDS-PAGE and HA-photocholesterol was visualized using the Typhoon FLA 9500 scanner (Excitation =555 nm; Emission =565 nm) in the native (unfixed) gel. In both experiments 50 µl (10%) of cell lysate was removed for western blotting to compare HA-wt and HA-YKLW4A expression levels. The density of HA bands of the western-blot and the fluorogram were analyzed with Image J software. The photo-crosslinking (fluorogram) to protein expression (western-blot) ratios were calculated for each mutant and experiment and results were normalized to HA wild-type = 100%. Results are show as mean plus/minus standard deviation.

### SDS-PAGE, Western-blot and HA2 antiserum

After sodium dodecyl sulfate-polyacrylamide gel electrophoresis (SDS-PAGE) using 12% polyacrylamide, gels were blotted onto polyvinylidene difluoride (PVDF) membrane (GE Healthcare). After blocking of membranes (blocking solution: 5% skim milk powder in PBS with 0.1% Tween-20 (PBST)) for 1h at room temperature, anti HA2 antibodies (diluted 1:2000 in blocking solution) were applied overnight at 4°C. After washing (3×10 min with PBST), horseradish peroxidase-coupled secondary antibody (anti-rabbit, Sigma-Aldrich, Taufkirchen, Germany, 1:5000) was applied for 1 hour at room temperature. After washing, signals were detected by chemiluminescence using the ECLplus reagent (Pierce/Thermo, Bonn, Germany) and a Fusion SL camera system (Peqlab, Erlangen, Germany). The density of bands was analyzed with Image J software.

Antisera against the HA2 subunit of FPV were generated in rabbits. Purified virus was subjected to SDS-PAGE and the Coomassie-stained HA2 band was cut from the gel and used for immunization.

### Determination of the cholesterol concentration

Cholesterol concentration in purified virus was determined using Amplex™ Red Cholesterol Assay Kit (Molecular Probes, Thermo Fisher) according to manufacturer’s instruction. Briefly, virus preparations (5 µl) purified from MDCK II cells with a 20-60% sucrose gradient were lysed in 1X reaction buffer (0.1 M potassium phosphate, pH 7.4, 50 mM NaCl, 5 mM cholic acid, 0.1% Triton X-100) and incubated with working solution (300 μM Amplex Red reagent, 2 U/ml Horseradish peroxidase, 2 U/ml cholesterol oxidase in 1X reaction buffer) at 37°C for 30 min in the dark. Cholesterol oxidase produces H_2_O_2_ that in the presence of horseradish peroxidase (HRP) reacts with the Amplex Red reagent in a 1:1 stoichiometry to produce highly fluorescent resorufin. Its fluorescence was measured using a microplate reader with an excitation wavelength of 555 nm and emission at 590 nm. We measured only cholesterol, not cholesterol esters, since virus samples were not treated with cholesterol esterase.

The protein concentration of the same virus preparations was measured with Roti-Quant universal kit (Carl Roth), which is based on the bicinchoninic acid (BCA) assay, except that PCA, a highly similar, but brighter molecule was used.

### Confocal microscopy

To study apical transport of HA with single or double mutations in the CCM (LA, YK2A and LW2A), polarized cells were infected with the respective viruses at an m.o.i of 1. Cells were fixed with 4% formaldehyde in PBS at 6 h post infection for 20 min and blocked with 3% BSA in PBS. Anti-HA2 antiserum (1:500) and monoclonal antibody against the basolateral marker ß-catenin (1:500) was then incubated with cells, followed by anti-rabbit secondary antibody coupled to Alexa Fluor 568 (red) and anti-mouse Alexa Fluor 488 (green), respectively, both at a dilution of 1:1000.

To study the apical transport of the HA mutant YKLW4A, 5×10^5^ MDCK II cells were seeded into 24 mm transwells one day before transfection using Lipofectamine 3000 Reagent (Invitrogen). 6 h post transfection, the upper chamber was changed to fresh DMEM supplemented with 2% FCS. The cells were cultured for 4 more days with changing medium every day. Cells were fixed with 4% formaldehyde in PBS for 20 min and permeabilized with 0.5% Triton X-100 for 5 min, followed by staining with primary antibody and secondary antibody as described above.

Cells were visualized with the VisiScope confocal FRAP System (VisiTron Systems GmbH), equipped with iXon Ultra 888 EMCCD camera, using 100X objective (1.45 NA) and illuminated via laser lines at 488 nm (Alex Fluor 488) and 561 nm (Alexa Fluor 568). Polarized cells were recorded in z-stacks with 0.5 µm increments and analyzed with Image J software.

### Membrane fusion assays

#### Hemolysis assay with virus particles

Culture supernatants of virus-infected MDCK II cells were cleared by low-speed centrifugation (2000 x g, 5 min) and were then adjusted with infection medium to a HA titer of 2^6^. 100 µl virus was added to 96-well plates with round bottom, mixed with 100 µl 2% chicken red blood cells (RBCs) in PBS and incubated at 4°C for 30 min. To pellet RBCs with bound virus samples were centrifuged at 250xg for 1 min. After removal of supernatant, the virus-RBC sediment was resuspended in 100 µl citric acid buffer (20 mM citric acid, 150 mM NaCl) adjusted with HCl to various mildly acidic pH values and incubated at 37°C for 1 h (or different time points between 0.5 and 4 h) to allow fusion between virus and RBCs. Then the plate was centrifuged at 250 g for 1 min and 50 µl of the supernatant was removed to determine the hemoglobin released from RBCs at a wavelength of 405 nm using a microplate reader.

To increase the cholesterol content in the viral membrane prior to hemolysis, 6 µl cholesterol stock (10 mM in chloroform: methanol (1:1; v:v)) was dried under nitrogen and resuspended in 150 µl methyl-ß-cyclodextrin (MßCD) solution (2 mM in aqua dest) which gives a molar ratio of 1:5 (0.4 mM/2mM). The mixture was shaken at 37°C overnight to load MßCD with cholesterol. Viruses grown in MDCK II cells were pelleted through a 20% sucrose cushion, resuspeded in 1X TNE buffer (10 mM Tris, 100 mM NaCl und 1 mM EDTA, pH 7.4) and adjusted to the same HA titer. 100 µl virus was incubated with 100 µl cholesterol-MßCD complex at room temperature for 30 min, centrifuged at 100000 g for 20 min to pellet the virus, which was then resuspended in 1xTNE buffer. 10 µl virus with an HA titer of 2^8^ loaded or not loaded with cholesterol was used for hemolysis assay.

#### R18 fluorescence dequenching assay with virus particles

Culture supernatants of virus-infected MDCK II cells were cleared by low-speed centrifugation (2000 x g, 5 min). Viruses were pelleted (100000 x g, 2 h) through a 20% sucrose cushion, resuspended in 1X TNE buffer (10 mM Tris, 100 mM NaCl, 1 mM EDTA, pH 7.4) and adjusted to a HA titer of 2^10^. For octadecyl rhodamine B chloride (R18) labeling, 50 µl virus was mixed with 0.5 µl 2 mM R18 (20 µM final concentration) and incubated on ice for 30 min in the dark. To remove unincorporated R18 samples were centrifuged at 100000 xg for 15 min at 4°C, labeled viruses were resuspended in 50 µl PBS and either used immediately or stored at −80°C.

To prepare erythrocyte ghosts, human RBCs were washed three times with PBS, lysed in ice-cold hypotonic buffer (4.7 mM Na_2_HPO^4^, 1.1 mM NaH_2_PO^4^, 1 mM EDTA, pH7.4) and again washed with PBS. For the fusion assay, 10 µl R18 labeled virus was incubated with 40 µl erythrocyte ghosts (1 mg/ml) at 4°C for 20 min. The virus-ghost mixture was then added to 1.96 ml prewarmed fusion buffer (150 mM NaCl, 10 mM Na-acetate x 3 H_2_O, pH 7.4) in a cuvette with a magnetic stir bar. Fluorescence intensity (Exitation: 560 nm, Emmission: 590 nm) was recorded at 37°C using the Cary Eclipse fluorescence spectrophotometer (Agilent Technologies). 100 s later when the fluorescence is steady, 7 µl citric acid (250 mM) was added to lower the pH to 5. 10 min after adding citric acid, 50 µl Triton X-100 (1% in aqua dest.) was added to the solution to achieve maximal dequenching. The fusion efficiency was calculated by the formula FDQ=100×(F(t)-F(0))/(F(max)-F(0)), with F(0) as the fluorescence intensity before adding citric acid, F(max) as the maximal dequenching intensity after adding Triton X-100 and F(t) as the fluorescence intensity at each time point. The equation used for curve fitting is f(x) = a* [1 – *e*^(−*kx*)^]

#### Double-labelled erythrocyte fusion assay with expressed HA

Human RBCs (1% in PBS) were double labeled with the lipidic dye R18 and the content marker calcein-AM (Molecular Probes, Life technologies). 20 µl of R18 (1 mg/ml in ethanol) was added to 1% RBC and incubated for 30min at room temperature in the dark. Samples were then washed twice with PBS and resuspended in 2 ml PBS. Calcein-AM (50 µg freshly dissolved in 10 µl DMSO + 40 µl PBS) was added to the mixture and incubated for 45 min at 37°C in the dark. The labelled RBCs were then washed with PBS five times and resuspended in 5 ml DPBS+. CHO cells in 6-well plate were transfected with HA-wt or HA-YKLW4A cloned into the pCAGGS vector. 24 h post transfection, cells were treated with 500 µl trypsin (5 µg/ml) plus neuraminidase (from *Clostridium perfringens*, 0.22 mg/ml, Sigma) in DPBS+ for 5 min at room temperature. The reaction was stopped by adding medium and cells were washed twice with DPBS+. 1 ml double-labeled RBCs were then added to HA-expressing cells and incubated at room temperature in the dark with gentle shaking. Unbound RBCs were removed by washing twice with DPBS+. DPBS+ adjusted with HCl to pH 5 was added to cells and incubated for 5min at 37°C. Acidic DPBS+ was replaced by DPBS+ adjusted to neutral pH and after incubation for 10 min cells were observed in an inverted fluorescence microscope (Zeiss, calcein channel: Excitation: band pass filter 470/540, Emission: BP 525/50; R18 channel: Excitation = BP 572/625 Emission =BP 629/662).

## RESULTS

### Mutation in the CCM decrease cross linking of HA to photocholesterol

The ectodomain of HA is connected by a nine amino acid long and flexible linker (that contains the cleavage sites for proteases used to remove the ectodomain from virus particles (39–41)) to the 26 amino acid long, α-helical transmembrane region (TMR) (42–44) and the eleven amino acid long cytoplasmic tail carrying three fatty acids attached to conserved cysteine residues (28, 45). Only HAs of the phylogenetic group 2 contain a CCM motif, comprising the conserved amino acids YK at the end of the linker and LW at the beginning of the TMR (Fig. 1A). A helical wheel plot revealed that the amino acids K, L and W are located on one, but Y on the other side of a helix suggesting that two helices of the trimeric HA molecule must contribute to binding of one cholesterol molecule (Fig. 1B). This distribution is thus similar to the amino acids that interact with cholesterol in 7TMR receptors, where the residues W/Y-I/V/L-K/R are on one helix, but another aromatic amino acid, either phenylalanine (F) or tyrosine (Y) on a second helix that bind cholesterol from the other side (32, 35, 36). The cholesterol binding site in the ß2-adrenergic receptor is located in the internal part of the transmembrane region and is completely embedded within the membrane (32), but other 7TMR receptors bind cholesterol to the outer part of the TMR or to amino acids that do not correspond to the CCM motif (35).

**Figure 1:**
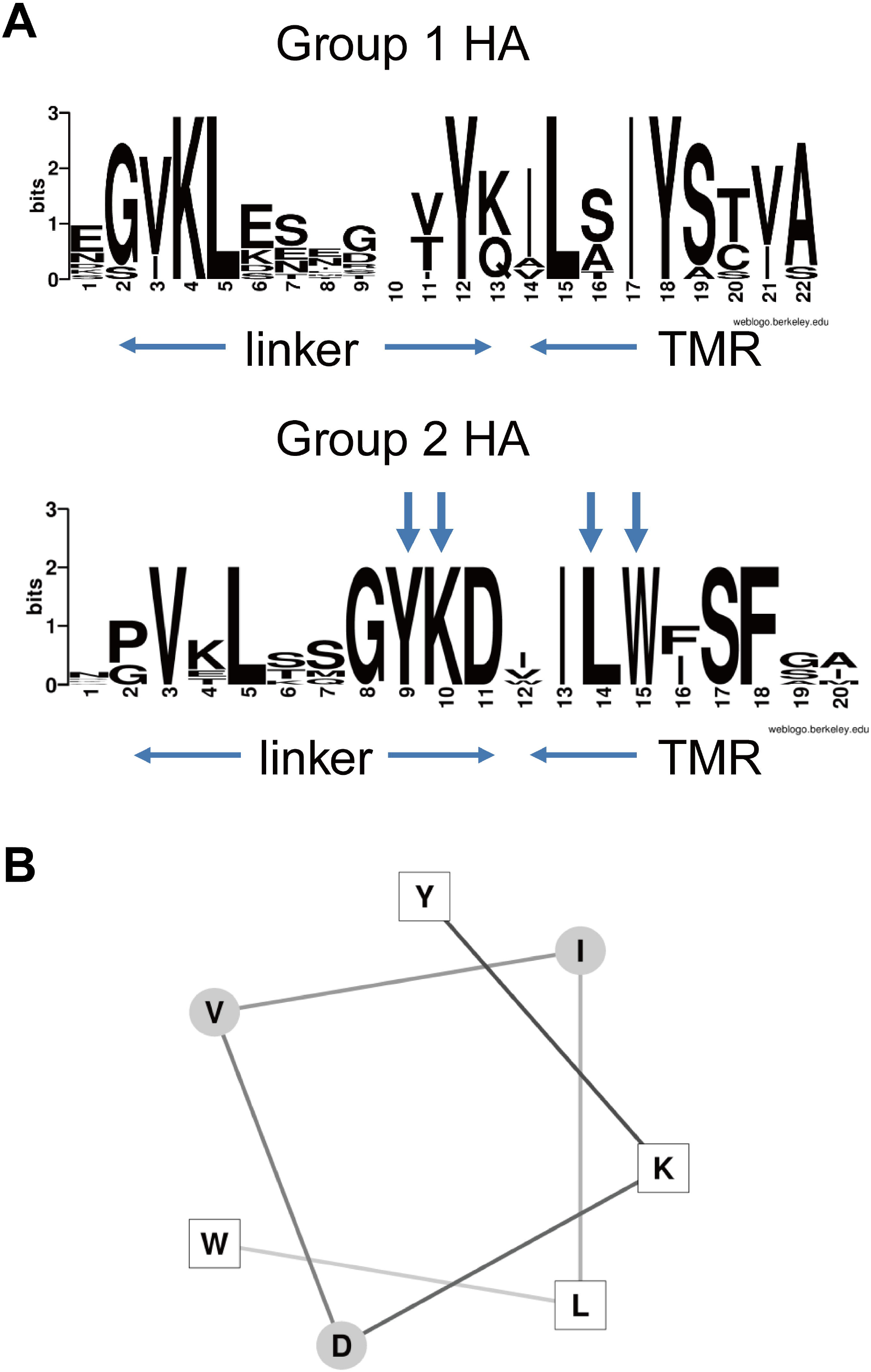
Cholesterol consensus motif (CCM) in HAs of the phylogenetic group 2. (A) Conservation of the CCM (amino acids YK…LW) in group 2, but not group 1 HAs. A consensus sequence for each HA subtype was assembled from each HA sequence present in the database (60). The consensus sequence of group 1 (H1, H2, H5, H6, H8, H9, H11, H12, H13, H16 subtypes) and group 2 HAs (H3, H4, H7, H10, H14, H15) was then used for the analysis by the WebLogo 3.3 server http://weblogo.threeplusone.com/create.cgi. The height of the stack indicates the sequence conservation, while the heights of each letter the relative frequency of an amino acid at that position. The start and end of the linker region and the start of the transmembrane region (TMR) are indicated by arrows. (B) Helical wheel projection (http://lbqp.unb.br/NetWheels/) of the sequence YKDVILW of H7 subtype HA. Amino acids forming the CCM are shown as white squares. Y, K, L are on one side of the helix; W is on the other side and thus must be located on another HA monomer of the trimeric spike to contribute to binding of the same cholesterol molecule, but this arrangement of TMR helices is speculative.

In our previous studies we reported that various mutations in the CCM severely retard transport of HA to the plasma membrane (31, 37), but whether the CCM interacts with cholesterol was not investigated. We performed experiments with a clickable–photocholesterol that contains a diazirine group at position 6 of the sterol ring (Fig. 2A). This moiety disintegrates upon uv-illumination into molecular nitrogen plus a highly reactive carbene-group that forms a covalent bond with amino acid side chains in close vicinity. To visualize cross-linked proteins, click-photocholesterol contains a terminal alkine group at the end of the side chain which can be “clicked” in a copper-catalyzed reaction under physiological conditions to the azido-fluorophore Cy3. A similar, but tritiated compound was used to demonstrate cholesterol-binding to synaptophysin (46) and another study showed that this photocholesterol is a faithful mimetic of authentic cholesterol (47). It is also more similar to genuine cholesterol than other photocholesterol probes used recently since it contains (besides the alkine group) no further alterations in cholesterol’s alkyl side chain (48). However, some of the diazirine groups might be photoactivated to other reactive species that have a longer half time than the carbene-group. They might then be cross-linked unspecifically to any proteins they encounter during diffusion through the membrane. Thus, photocrosslinking is a qualitative rather than a quantitative measure of the cholesterol affinity of a protein. Nevertheless, all available compounds label only a few specific proteins out of all cellular membrane proteins (46, 48) indicating that they are suitable tools to identify proteins that strongly (but non-covalently) interact with cholesterol.

**Figure 2:**
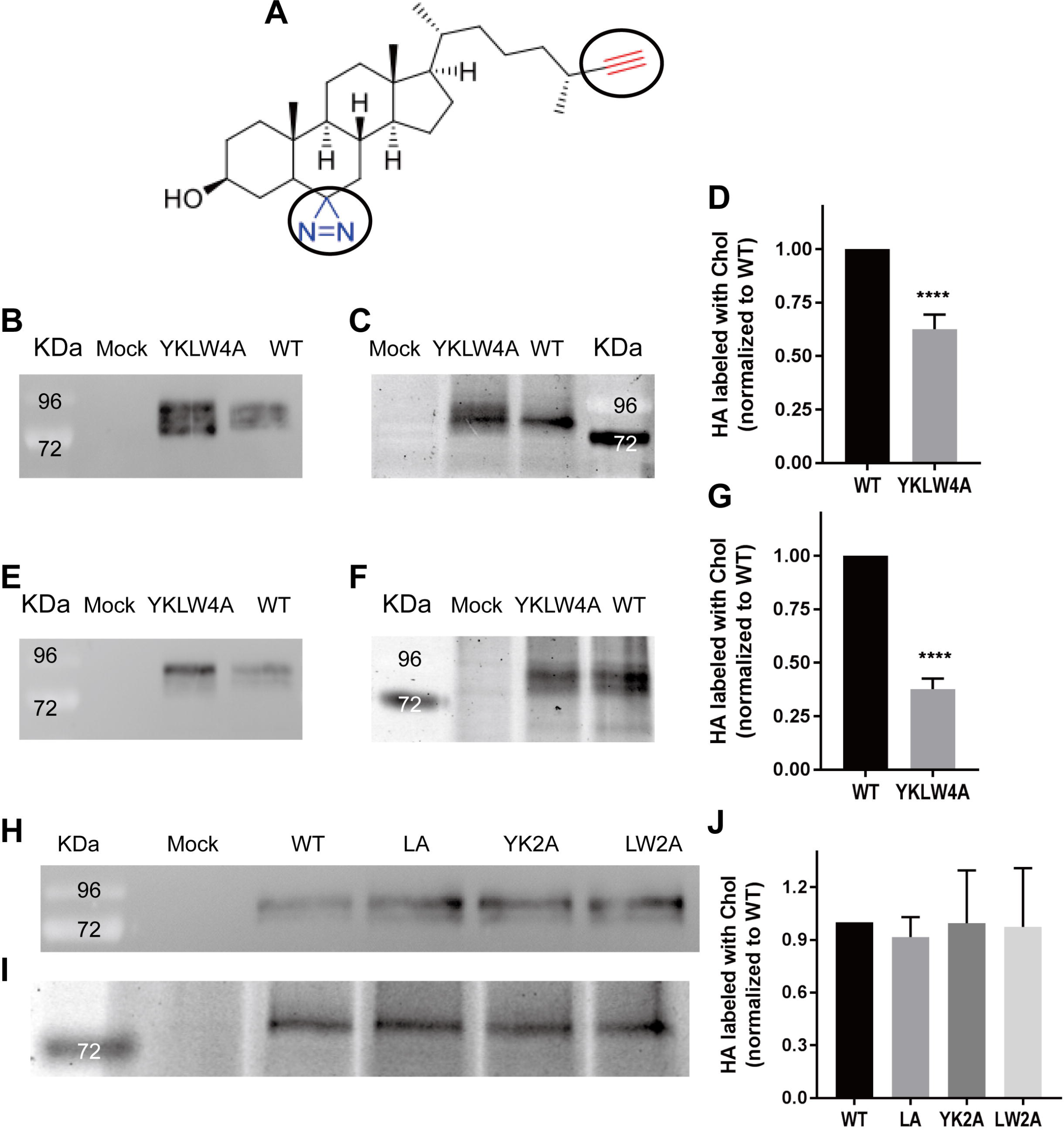
Photocholesterol labeling of HA wt and HA with mutations in the CCM. (A) Formula of click-photocholesterol (6’-Azi-25-ethinyl-27-norcholestan-3ß-ol). The functional groups diazirine (blue) and azide (red) are encircled. (B-D) Labeling of CHO-cells expressing HA wt or HA YKLW4A. After labeling for 16 hours, cells were exposed to uv-light and lysed with 1% Triton. 10% of the lysate was subjected to SDS-PAGE followed by western-blotting with HA2 specific-antiserum to monitor expression levels of HA (B). 90% of the lysate was immunoprecipitated with the same antiserum, HA-photocholesterol was clicked to Cy3 fluorophore, samples were subjected to reducing SDS-PAGE and the fluorescence in the gel was scanned (C). Quantification: Band intensities of (B) and (C) of this and five other experiments were determined and normalized to HA wt = 100%. The mean (58%) ± standard deviation (±13%) is shown (D). (E-G) Labeling of immunoprecipitated HA wt or HA YKLW4A. 24 hours after transfection CHO cells expressing HA wt or HA YKLW4A were lysed. 10% of the lysate was subjected to SDS-PAGE followed by western-blotting (E). 90% of the lysate was immunoprecipitated, washed antibody-HA complexes were incubated with photochlesterol, uv-irradiated and clicked to Cy3 fluorophore prior to reducing SDS-PAGE and fluorescence scanning (F). Quantification: Band intensities of (E) and (F) of this and three other experiments were determined and normalized to HA wt = 100%. The mean (38%) ± standard deviation (±5%) is shown (G). The asterisks indicate statistically significant differences (****P < 0.0001) between wt and the mutant according to a Student’s t-test. Mock: untransfected cells. kDa: Molecular weight markers. HA YKLW4A was in each experiment (B-E) expressed at higher levels as HA wt. (H+I) Labeling of CHO-cells expressing HA wt, HA LA, HA YK2A and HA LW2A. Experiment was performed as described in (B) and (C). Quantification: Band intensities of (H)) and (I) of this and two other experiments were determined and normalized to HA wt = 100%. The mean ± standard deviation is shown (J). HA LA 0.92±0.11, YK2A 0.99±0.30 and LW2A 0.97±0.34 relative to HA wt.

We expressed H7 subtype HA from a variant of fowl plague virus (FPV*) having a monobasic cleavage site, both the wild-type protein and a mutant where the four amino acids forming the CCM were replaced by alanine (HA YKLW4A). In the first experiments transfected CHO cells were labeled with click-photocholesterol for 16 hours and subsequently uv-irradiated for 10 minutes. Cells were then lysed, one aliquot was subjected to western-blotting with HA2 specific antibodies, the other aliquot to immunoprecipitation using the same antibodies and click-chemistry. The resulting fluorescence scan showed incorporation of photocholesterol into both HA wt and HA YKLW4A in approximately similar amounts (Fig. 2C). However, the western blot revealed that the expression level of HA YKLW4A is significantly higher (Fig. 2B). Quantification of fluorescence intensities and normalizing them to the expression level showed that incorporation of photocholesterol into HA YKLW4A was reduced to 58% (±13%, mean of six transfections, Fig. 2D).

Since mutations in the CCM decrease association of HA with nanodomains (27, 37) one might argue that the diminished labeling of HA YKLW4A might be due to its compartmentalization into cholesterol-depleted membrane domains. Consequently, less cholesterol (and hence photocholesterol) is present in the vicinity of HA YKLW4A and thus available to label the protein by random interactions. To exclude such an unspecific effect, we first immunoprecipitated HA wt and HA YKLW4A from cell lysates and then performed photo-crosslinking and click-chemistry on the purified HA-antibody complex (Fig. 2E+F). Nevertheless, a similar result was obtained; incorporation of photocholesterol into HA YKLW4A was even more reduced relative to HA wt (38 ±5%, mean of four transfections, Fig. 2G).

To determine whether partial exchange of the CCM has an effect on photo-crosslinking we created HA double mutants HA LW2A and HA YK2A where two consecutive amino acids located at the end of the TMR and in the linker region, respectively were exchanged by alanine. In HA LA the leucine in the TMR (which of all single mutants had the strongest effect on intracellular transport of HA (37)) was substituted by alanine. Cells expressing the three HA mutants were labeled and analyzed as before but no significant reduction of incorporation of photocholesterol was detected (Fig. 2H-J). Thus, in order to reduce photo-crosslinking of HA the whole CCM must be exchanged suggesting that the residues act synergistically with cholesterol.

### Mutations in the CCM of HA affect virus replication

All our previous experiments about the CCM of HA were performed with expressed protein (37). To investigate whether cholesterol binding to HA affects virus replication, we created the described mutations in the CCM of HA in the context of the viral genome. We used a variant of fowl plague virus (A/FPV/Rostock/34, H7N1), termed FPV* which contains a monobasic cleavage site in HA and thus requires trypsin for growth in cell culture (38). The amino acid exchanges were generated by at least three nucleotide substitutions to exclude that mutant viruses revert back to wild type.

The mutant HA plasmid together with plasmids encoding the other viral proteins were transfected into HEK 293T cells, the supernatant was used to infect MDCK II cells and release of virus particles was assessed by HA assays. In three independent transfections we never rescued virus particles for FPV* YKLW4A, whereas wild-type virus and the other three mutants done in parallel exhibit HA titers of 2^5^ −2^6^. From the rescued mutants a virus stock was generated in MDCK II cells and sequencing of the HA gene showed that the desired mutations were still present (data not shown).

To compare the replication kinetics of the viruses, MDCK II cells were infected with FPV* wt or with the mutants at an m.o.i. of 0.0005 (based on TCID50 titer), supernatants were collected at various time points post infection and virus titers were assessed by HA and TCID 50-assay (Fig. 3A+B). The growth curve revealed a statistically significant decrease in the TCID 50 titer for FPV* LW2A (1.5 logs, ∼95% reduction, mean of 3 experiments) at 36 and 48 hours post infection. Titers of FPV* YK2A and FPV* LA were also somewhat decreased relative to FPV* wt.

**Fig. 3:**
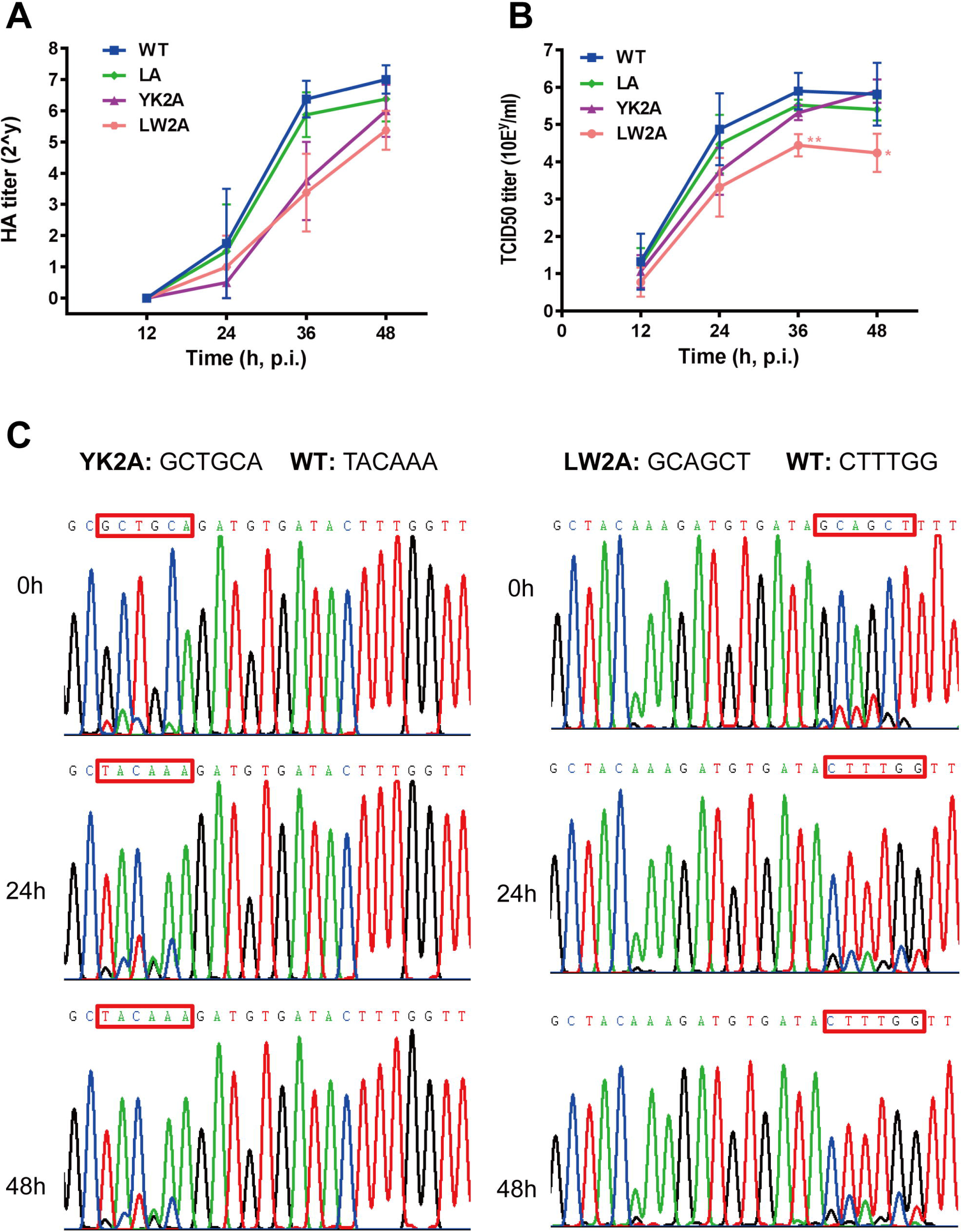
Effect of mutations in the CCM on virus replication. (A+B) Growth curves of FPV* wt and FPV* with the indicated mutations in the CCM of HA. MDCK II cells were infected with virus at an m.o.i. of 0.0005. Culture supernatants were harvested at the indicated times and tested with HA-assay (A) or TCID50 (B). Experiments were carried out in triplicate and are displayed as means±standard deviation. Asterisk indicate statistically significant differences (*P < 0.05, **P < 0.01) between wt and the mutant LW2A according to a Student’s t-test. (C) Competitive growth of FPV* wt and FPV* mutants. Sequencing chromatograms of cDNA of wild-type and mutant FPV*. MDCK II cells were infected (total moi: 0.0005) with FPV* wt and FPV* YK2A (left) or FPV* LW2A (right) mixed at ratio of 1:5. Supernatants were collected before or at the indicated times after infection, the viral RNA was isolated and subjected to rtPCR and sequencing. The nucleotide sequences of wt and mutant HA are listed above the chromatogram.

We next asked whether mutating the CCM of HA might reduce the competitive fitness of the virus. To test this, we mixed FPV* YK2A or FPV* LW2A with FPV* wt at a ratio of 5:1, co-infected MDCK II cells (total moi of 0.0005), extracted viral RNA from cellular supernatants, either before or at 24 and 48 hours after infection, amplified the relevant part of the HA gene with rtPCR and analyzed it by sequencing. Fig. 3C shows the sequencing chromatograms for the region of interest in the HA gene. Both wild-type and mutant viruses were detected at all time points, reflected by superimposed peaks for the respective nucleotide bases at the mutation site. Due to the higher number of infectious mutant viruses in the inoculum the mutant sequence is predominant before infection, but particles released from cells after 24 and 48 hours contain mainly the wild type sequence. Although the differences in the peak heights in the chromatograms should not be interpreted in a precise quantitative manner, it is obvious that the wild type virus rapidly outgrows mutant virus with two exchanges in the CCM within a few replication cycles.

In sum, we conclude that the CCM is essential for virus replication. Exchanging all four amino acids of the motif prevented generation of infectious virions and exchanging two of them reduced virus titers and their competitive fitness.

### Mutations in the CCM do not cause mistargeting of HA in polarized cells

In polarized MDCK cells HA is transported to the apical membrane, the viral budding site (49). Since signals for transport are located in the TMR of HA (50–52) we analyzed whether mutations in the CCM disturbed polarized budding of virus particles. MDCK cells grown on transwell filters were infected with FPV* wt and FPV* mutant virus at low moi, aliquots of the supernatant were removed from the apical and basolateral chamber at various time points and virus titers were determined using HA-assays. The growth curve plotted for virus particles released from the apical membrane shows again a reduction in virus titers for all FPV* mutants, statistically significant for FPV* LW2A. However, no virus was released for any of the mutants from the basolateral membrane (Fig. 4A). Since Influenza virus might bud from the apical membrane even if HA is redirected to the basolateral membrane (53) we also determined the localization of HA LA, HA LW2A and HA YK2A in virus-infected and of HA YKLW4A in transfected cells using confocal microscopy. However, wt HA and all mutants are exclusively located at the apical membrane, the fluorescence derived from anti-HA antibodies does not overlap with the fluorescence emitted by the basolateral membrane marker (Fig. 4B+C). We conclude that mutations in the CCM do not affect targeting of HA to the apical membrane.

**Fig. 4.**
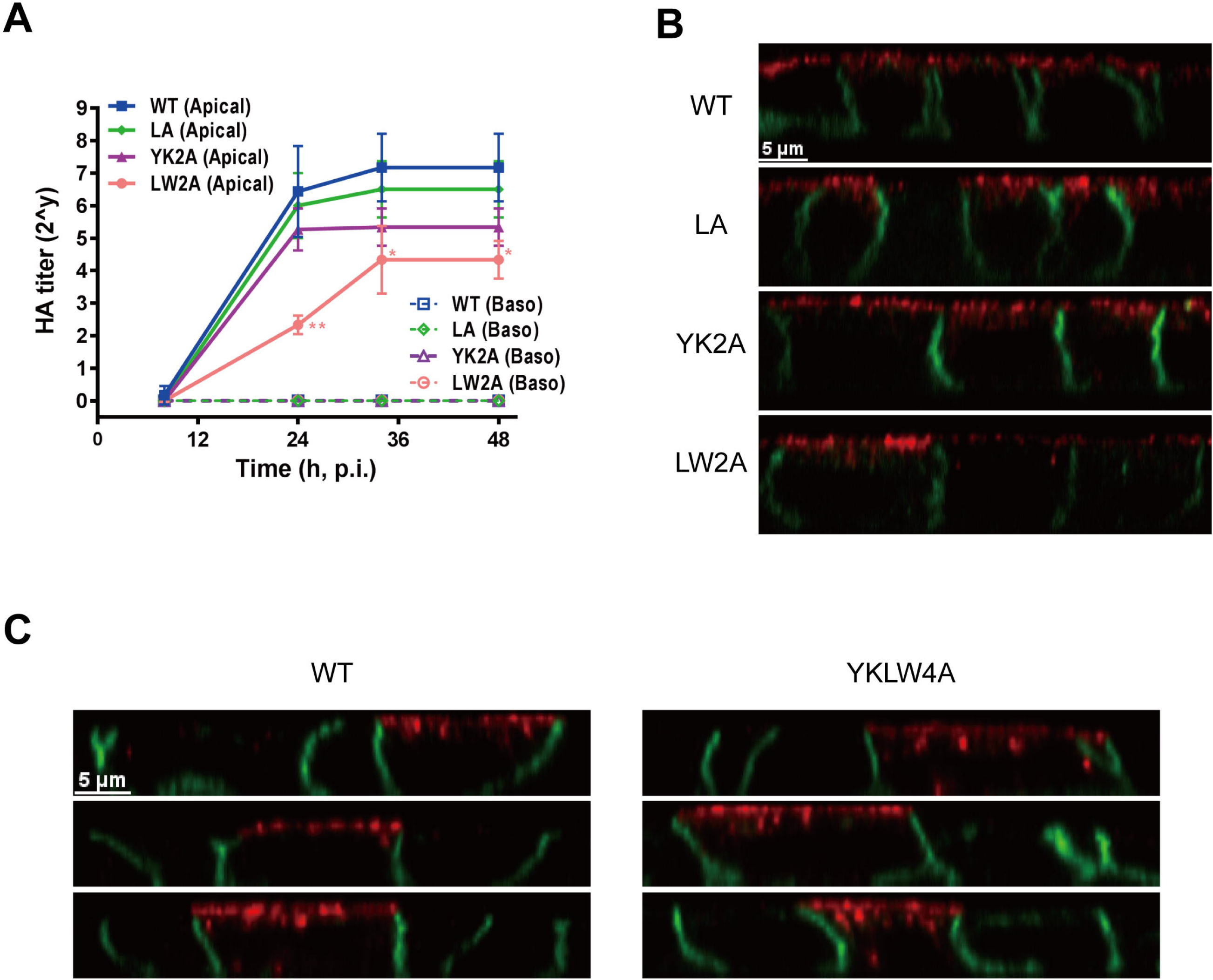
Effect of mutations in the CCM on apical virus budding and transport of HA in polarized cells. (A) Virus budding in polarized MDCK II cells. Polarized MDCK II cells were infected with viruses at a m.o.i. of 0.001. Culture medium from both the upper (apical) and lower (baso) chamber was harvested at 8h, 24h, 34h, and 48h post infection and virus titers were determined by HA-assays. Asterisk indicate statistically significant differences (*P < 0.05, **P < 0.01) between wt and the mutant LW2A according to a Student’s t-test. (B) HA mutant LA, YK2A and LW2A transport in MDCK II cells. Polarized MDCK II cells were infected with the respective viruses at a m.o.i. of 1. 6h post infection, cells were fixed and stained with anti-HA2 antiserum and anti-ß-catenin antibody (basolateral marker), followed by secondary antibody coupled to Alexa Fluor 568 (red for HA) and Alexa Fluor 488 (green for catenin), respectively. (C) HA mutant YKLW4A transport in polarized MDCK II cells. MDCK II cells were transfected with HA mutant YKLW4A and HA wt one day after they were seeded into 24 mm transwell filter membranes. 4 days post transfection; cells were fixed and permeablized, and stained with primary and secondary antibody as described in (B). Z-sections with 0.5 µm increments from polarized cells are shown in (B) and (C). Staining of intracellular structures is visible in (C) especially for YKLW4A since the cells were permeabilized.

### Mutations in the CCM reduce incorporation of HA and cholesterol into virions

In principle, the decrease in virus replication could be due to compromised cell entry of mutant virus and/or a defect in virus assembly and budding. We have previously shown that mutations in the CCM decreases HA’s cell surface exposure and association with membrane rafts in transfected cells (37). Since these defects might affect incorporation of HA into budding virions we purified FPV* wt and FPV* LW2A virus particles with sucrose gradient centrifugation from embryonated eggs and analyzed their protein composition by SDS-PAGE and Coomassie staining (Fig. 5A). Densitometry of viral protein bands and calculation of the viral protein ratios (Fig. 5B) revealed for FPV* LW2A reduced amounts of HA relative to NP (85%, normalized to FPV* wt) and to M1 (82%). If virus particles were purified from MDCK II cells (Fig. 5C), the reduction in HA incorporation was even more pronounced; the HA/M1 ratio is reduced to 59% and the HA/NP ratio to 68%. Likewise, the other two mutants also exhibit a (albeit less distinct, 82-90%) reduction in the HA content (Fig. 5D).

**Fig. 5:**
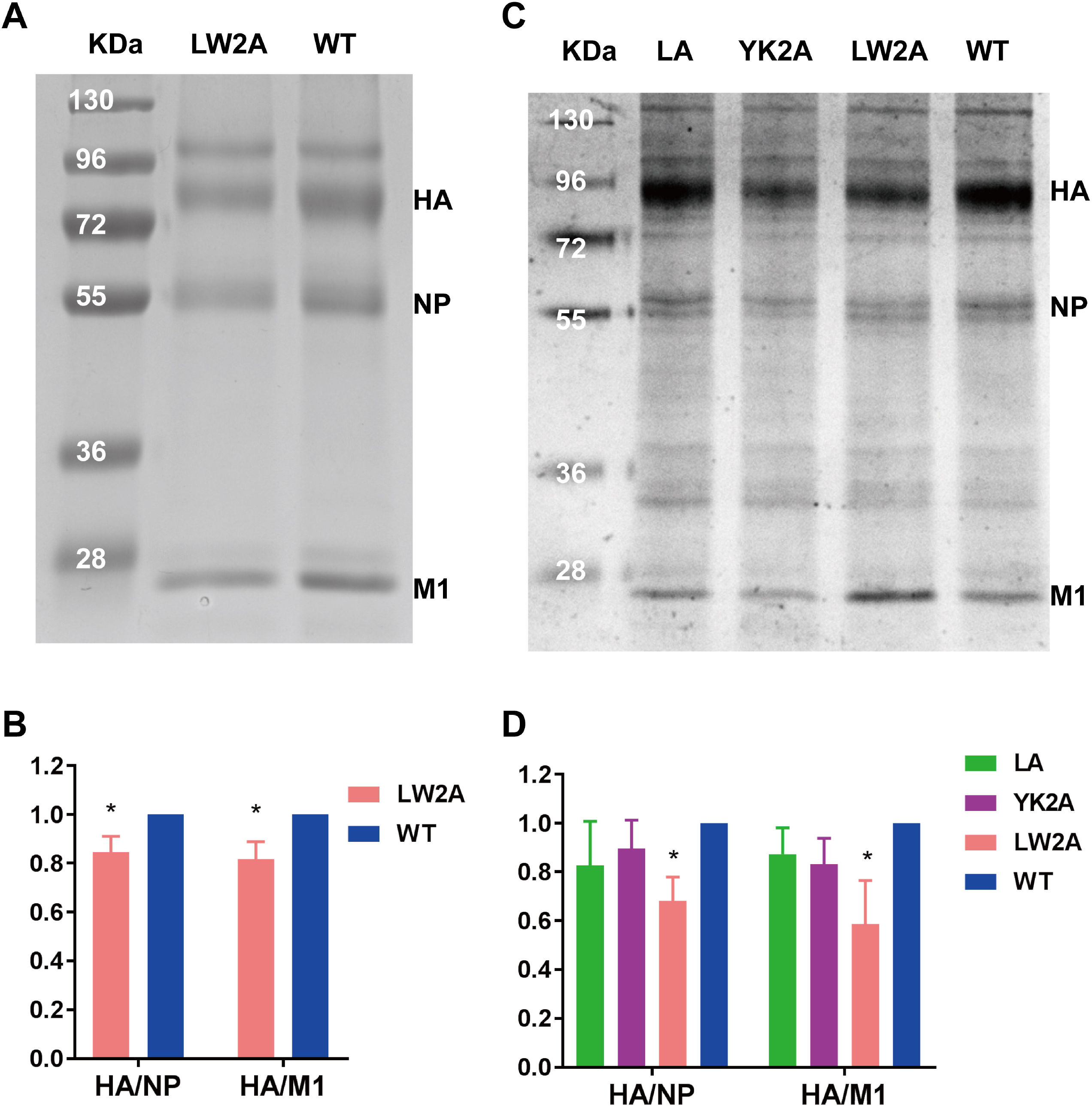
Effect of mutations in the CCM on HA incorporation into virus particles. (A+C) Protein composition of FPV* particles purified from embryonated eggs (A) and MDCK II cells (C). Viruses were purified using a sucrose gradient (20%-60%) and subjected to non-reducing SDS-PAGE and Coomassie staining. The position of the major viral proteins is indicated on the right-hand side, and molecular mass markers (kDa) are shown on the left-hand side. (B+D) Quantification of the relative protein composition. Density of HA, NP and M1 bands was determined and the HA/NP and HA/M1 ratios were calculated and normalized to wild type. Results from three virus preparations are displayed as means ±standard deviation. Asterisk (*) indicate statistically significant differences (*P < 0.05) between wt and the mutant LW2A according to a Student’s t-test.

Since mutations in the CCM reduce association of HA with membrane nanodomains (27, 37) and since FPV wt buds through cholesterol-enriched domains (5) we asked whether FPV* mutants possess less cholesterol in their membrane. We therefore determined the cholesterol concentration of the same three virus preparations from MDCK II cells and divided it by the protein concentration. For FPV* wt we determined a mean of 220 nM cholesterol per ng/µl total protein, but considerable variations between individual preparations was observed. Nevertheless, the cholesterol content of each FPV* mutant was lower in every (except one) virus preparation relative to FPV* wt, which was purified from sister cultures in parallel. The only exception was FPV* LA for which a slightly higher amount of cholesterol was determined in one, but not in the other two preparations (see Fig. 6A for details on individual experiments).

**Fig. 6:**
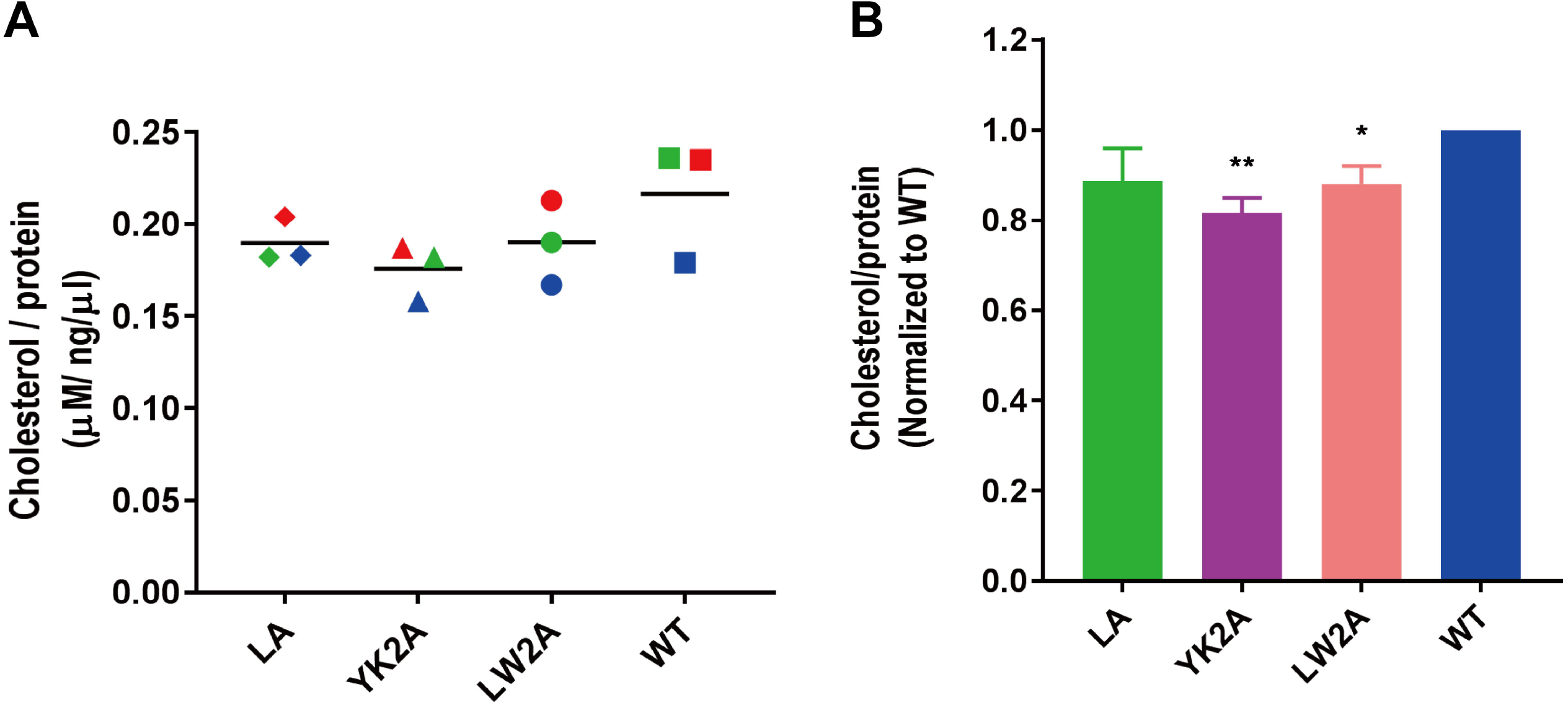
Effect of mutations in the CCM on cholesterol content and virus morphology. (A) Cholesterol content of FPV* wild-type (wt) and FPV* mutant particles sucrose-purified from MDCK II cells. Dots indicate the cholesterol concentration (µM) divided by the protein concentration (ng/µl) for each of three virus preparations; the horizontal bar is the mean. Viruses prepared from sister cultures and analyzed in parallel are indicated with the same color. (B) Normalized cholesterol content (wt=100%) of FPV* mutant particles. Asterisk (*) indicate statistically significant differences (*P < 0.05, **P < 0.01) between wt and the mutant YK2A or mutant LW2A according to a Student’s t-test.

Normalizing the cholesterol content for each virus preparation (wt =100%) exhibit a reduction to 89% in FPV* LA, to 88% in FPV LW2A and to 82% in FPV YK2A (Fig. 6B). Assuming that our FPV* wt preparations contain 52 mol% of cholesterol in relation to all other envelope lipids as determined by quantitative mass spectrometry for FPV particles grown in the same cell type (5), one can calculate that the cholesterol content in mutant particles is reduced to 46% in FPV* LA and FPV* LW2A and to 43% in FPV* YK2A. This corresponds quite exactly to the cholesterol content (45%) determined for the apical membrane of polarized MDCK II cells (5). Thus, our results are consistent with the concept that viruses with mutations in the CCM bud not through (cholesterol-enriched) membrane nanodomains but through the bulk phase of the plasma membrane (26).

Influenza A virus mutants with defects in virus assembly and budding release particles with aberrant morphology (54). We therefore investigated the virus preparations also by negative stain electron microscopy, but no differences in the morphology or size of intact virus particles were obvious (data not shown). Whether, the lower cholesterol content affects the density of HA spikes in virus particles requires investigations with more sophisticated methods (25).

### Mutations in the CCM decrease the hemolytic activity of HA

Next, we investigated whether mutations in the CCM affect cell entry of viruses via HA-mediated membrane fusion. Since the extent of membrane fusion increases with the HA-concentration and since the HA amount is reduced in mutant virus preparations (Fig. 5A-D) we first used hemolysis assays. This allows adjusting the amount of virus by means of its HA-titer and to record membrane fusion quantitatively by measuring the release of hemoglobin.

In the first set of experiments we compared the pH dependence of hemolysis. Wild type and mutant virus with an HA-titer of 2^6^ were adsorbed to chicken erythrocytes, the pH was adjusted to mildly acidic pH values between 5 and 7, erythrocytes with bound virus were incubated for 60 min at 37°C and hemoglobin release was determined spectroscopically. Beginning with pH 5.7 FPV* wt causes hemoglobin release; the amount increased linearly with decreasing pH values (Fig. 7A) which is in agreement with published data on the fusion activity of the closely related H7 Weybridge strain (55). All mutant viruses also start to cause hemolysis at pH 5.7, but the amount of released hemoglobin is reduced to ∼ 30% with each virus and at all pH values. We also compared the kinetics of hemolysis by incubating pH 5 activated viruses for up to four hours with erythrocytes. The amount of released hemoglobin increases linearly with time for all viruses, but the slope of the line is much steeper with FPV* wt. At every time point the amount of released hemoglobin is clearly lower when using mutant virus particles (Fig. 7B).

**Fig. 7:**
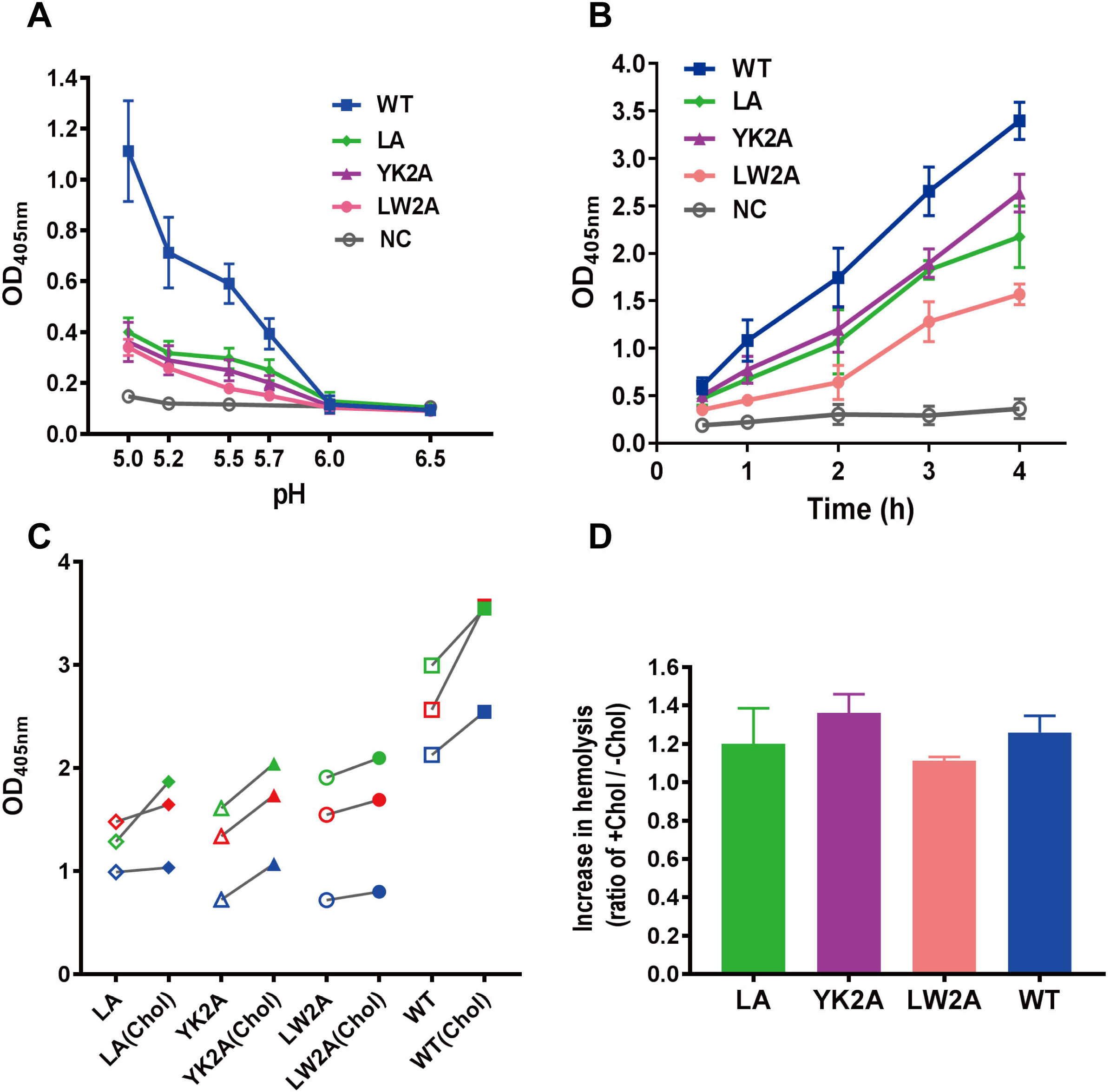
Effect of mutations in the CCM on hemolysis. (A) pH dependence: FPV* wt and FPV* mutant particles were adjusted to an HA-titer of 2^6^, adsorbed to chicken erythrocytes and pelleted. Samples were adjusted to the indicated pH values and incubated for 60 min at 37°C. Released hemoglobin (OD 405,) is plotted against the virus titer. Results are depicted as means±standard deviation from three experiments. NC: negative control: Incubation of RBCs without virus. (B) time dependence: Hemolysis was initiated with pH 5 treatment and aliquots were removed after 0.5, 1, 2, 3 and 4 hours. Results are depicted as means±standard deviation from three experiments. (C+D): Comparison of hemolytic activity before and after loading of FPV* wt and FPV* mutant virus particles with cholesterol. Hemolysis was performed after acidification to pH 5 for 60 min at 37°C with viruses adjusted to an HA-titer of 2^8^. C: OD measurements of individual experiments with virus before (open symbols) and after (closed symbols) loading with cholesterol. Viruses prepared from sister cell cultures, loaded with cholesterol and measured in parallel are indicated with the same color. Note that in each experiment the hemolytic activity of mutant virus particles after loading with cholesterol is lower than the corresponding wild type viruses before cholesterol loading. D: Normalized increase in hemolytic activity relative to untreated virus particles (=1) from the same virus preparation. Results of three experiments are shown as mean±standard deviation.

Withdrawal of cholesterol from the viral membrane negatively and addition of cholesterol positively affect HA’s membrane fusion activity (22, 23, 56, 57). In principle, the defect in the fusion activity of mutant viruses might be either due to altered biophysical properties of the viral membrane caused by their reduced cholesterol content or due to a more local or intrinsic functional defect in the HA molecule. To distinguish between both possibilities, we replenished the viral membrane with cholesterol by incubation of virus particles with cyclodextrin fully loaded with this lipid. Cholesterol measurements of virus preparations before and after incubation with cyclodextrin revealed for FPV* wt and all mutant viruses a cholesterol increase of ∼20-35%; mutant viruses have now a cholesterol content similar to or slightly higher than FPV* wt before cholesterol addition. Hemolysis assays at pH5 showed an increase in hemoglobin release for both wild-type and mutant viruses by 10% to 35% after cholesterol loading (Fig. 7D) confirming the beneficial effect of cholesterol on membrane fusion. However, the hemolysis activity of cholesterol-enriched mutant viruses was in each experiment lower compared to untreated wild-type virus particles (Fig. 7C) indicating that the defect in membrane fusion persists even if the mutant particles have a cholesterol content like wild-type viruses.

### Mutations in the CCM decrease the hemifusion activity of HA

A defect in hemolysis does not reveal which step in membrane fusion is affected, since release of hemoglobin requires lipid mixing as well as opening and widening of a fusion pore. In some HA mutants both events are uncoupled, for example HA with a glycolipid-anchor instead of the transmembrane domain (GPI-HA), causes hemifusion, but not full fusion (17). We were therefore interested whether or not viruses with mutations in the CCM of HA exhibit a defect in hemifusion. We employed the R18 dequenching assay that allows monitoring lipid mixing between erythrocyte ghosts and virus particles. The lipophilic fluorophore octadecylrhodamine (R18) is integrated into the viral envelope at self-quenching concentrations. Upon binding of washed virus particles (adjusted to an HA-titer of 2^10^) to ghosts and activation of HA’s fusion activity by low pH treatment, viral and ghost lipids begin to mix. This causes dequenching of R18 and the resulting fluorescence increase is recorded in a fluorescence spectrometer. Once viral membrane fusion is completed, detergent is added that causes complete lysis of membranes. The maximal dequenching of R18’s fluorescence is used to calculate the fusion activity at each time point.

Fig. 8A shows the mean relative fluorescence intensity of four fusion reactions plotted against the time course. After acidification to pH5 all FPV* viruses exhibit a rapid increase in the fluorescence intensity, which reaches a plateau after ∼ 2min. FPV* wt exhibits a maximal fusion activity of 20%, which is clearly decreased in the mutants. Normalizing the extent of fusion (FPV* wt = 100%) shows a reduction to 75% for FPV* LA and to ∼ 50% for the double mutants FPV* LW2A and YK2A (Fig. 8B). Furthermore, fitting the curves revealed also differences in the fusion kinetics. 36 secs is the half time for maximal fusion calculated for FPV* wt and FPV* LA, but this is extended to 57 secs for FPV* LW2A and FPV* YK2A. Since a similar result was obtained if the R18 assay was performed at pH 5.5 (except that the half times were longer for all viruses, data not shown), we conclude that mutations in the CCM of HA affect both the kinetics and the extent of lipid mixing.

**Fig. 8:**
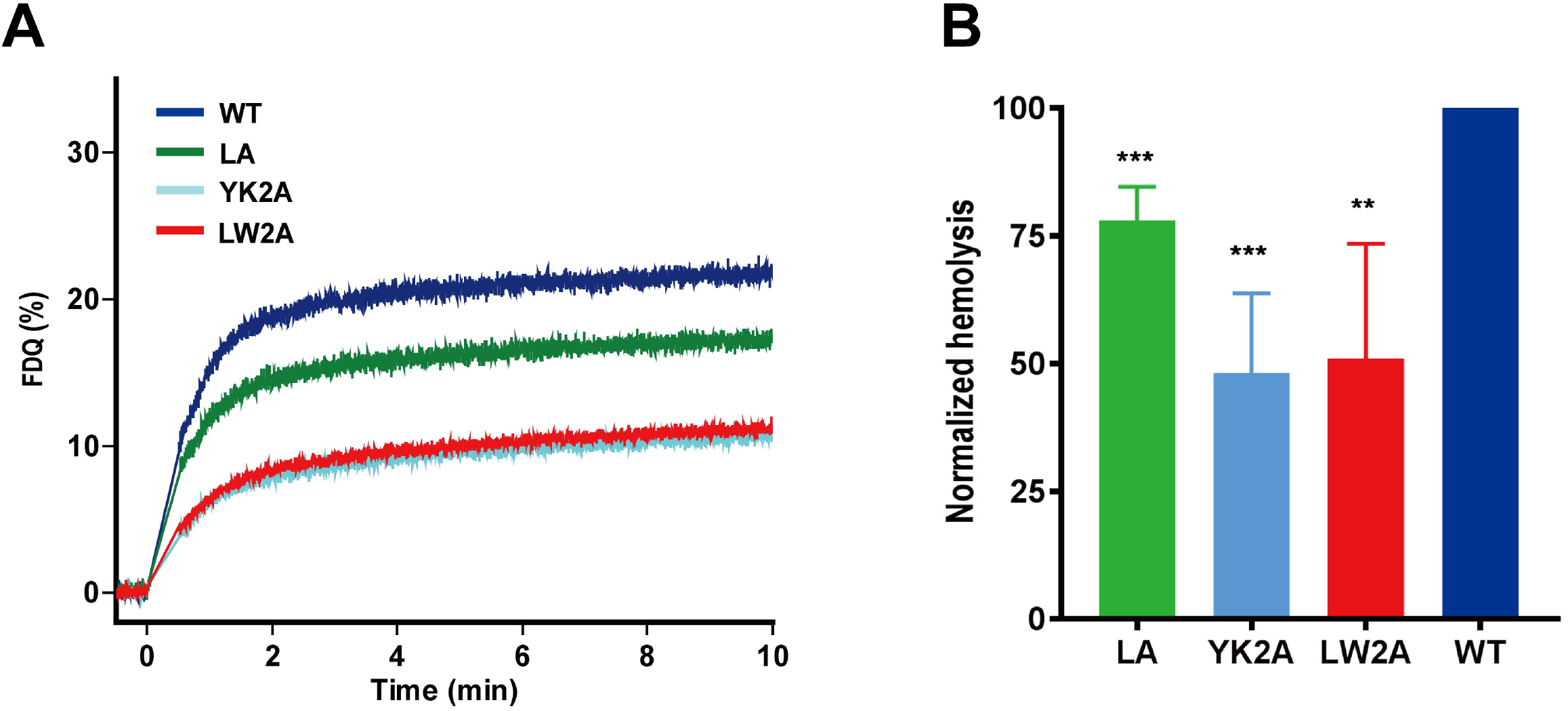
Effect of mutations in the CCM on hemifusion. (A) Fluorescence dequenching assay using erythrocyte ghosts labelled in the membrane with the self-quenching lipophilic fluorophore R18. FPV* wt and mutant viruses were adjusted to an HA-titer of 2^10^, adsorbed to R18-labeled ghosts and the fusion reaction was started by adjusting the pH to 5. The graph shows the mean of four experiments with two virus preparations. Relative fluorescence dequenching (FDQ, dequenching with Triton-X-100 = 100%) is plotted against the time (min). (B) Relative extent of fusion (FDQ after 10 min and normalized to wt =100%) is shown as the mean±standard deviation of the four experiments. Asterisk indicate statistically significant differences (*P < 0.05, **P < 0.01, ***P < 0.001) between wt and the mutants according to a Student’s t-test. Fitting the curves revealed also a delay in the half time for fusion for the double mutants. The following data were calculated: wt=0.59±0.05 min, LA=0.6±0.1; YK2A= 0.9±0.18; LW2A=0.95±0.1.

### HA with a complete exchange of the CCM has a defect in hemifusion

Finally, we asked whether the HA mutant with a complete exchange of the CCM also exhibits a defect in membrane fusion. Since no virus particles could be rescued for that mutant we had to rely on a cell-based fusion assay. As target we used double-labeled erythrocytes that contain R18 in their membrane and the soluble fluorophore calcein in their interior. After proteolytic cleavage and acid treatment of HA, R18 diffuses into the cellular plasma membrane whereas calcein stains the cytoplasm which can both be monitored in the fluorescence microscope.

Cells expressing HA YKLW4A clearly show hemifusion (cells in the upper left quadrant of Fig. 9A) and also full fusion (some cells in the right half of the figure) with erythrocytes. However, the number of fusion events is clearly reduced compared to HA wt, see Fig. 9B for an image of cells expressing HA YKLW4A or HA wt at lower magnification. One factor contributing to the lower fusion efficiency might be the retarded intracellular transport and reduced surface expression of HA YKLW4A (37). However, the number of bound erythrocytes per transfected cell culture plate is not obviously reduced with cells that express HA YKLW4A (micrograph in Fig. 9B) and cells transfected with HA YKLW4A show similar hemadsorption activity as cells transfected with HA wt (not shown).

**Fig. 9:**
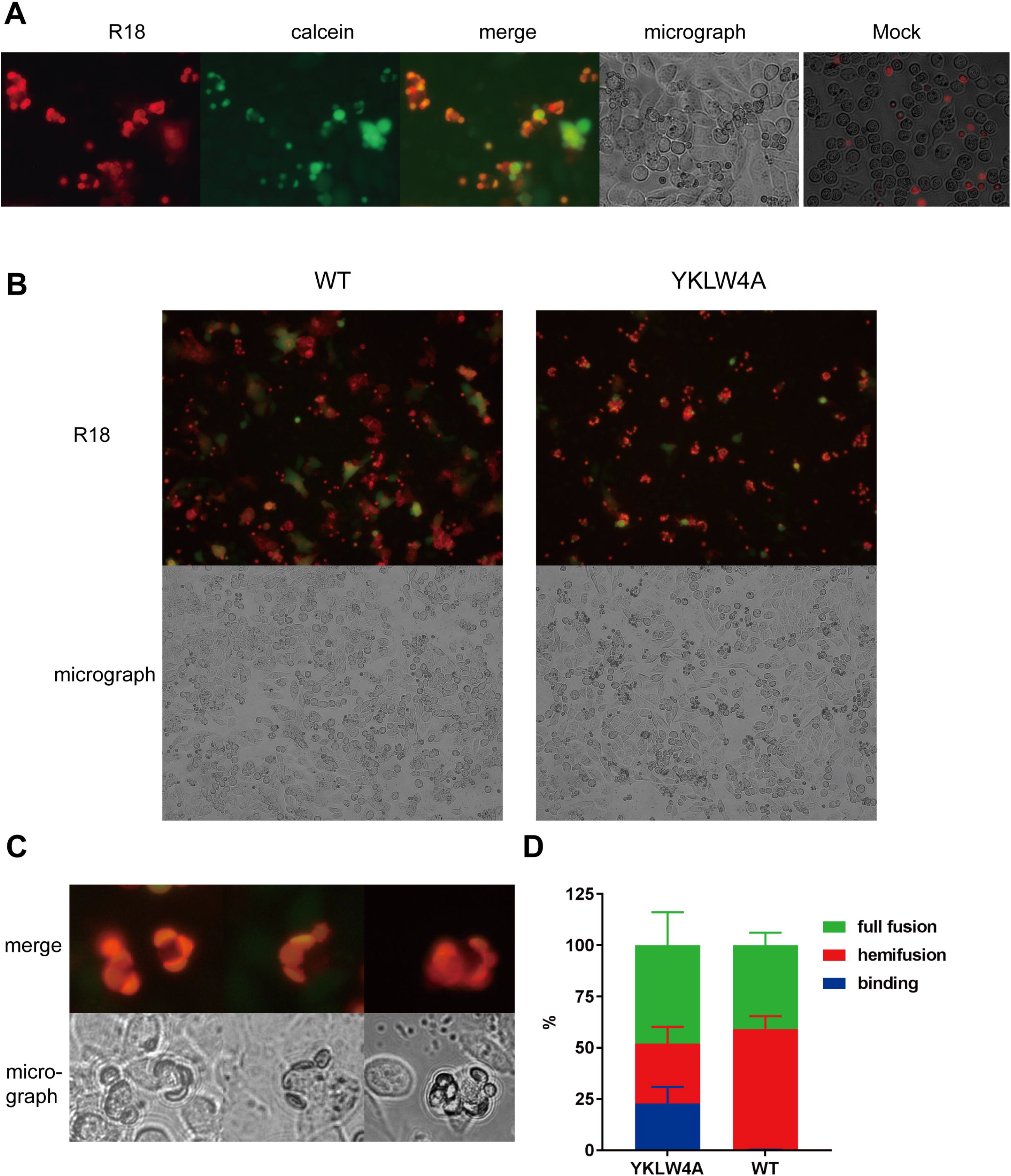
Effect of complete exchange of the CCM on fusion of cells to erythrocytes. Fusion of CHO cells expressing HA YKLW4A or HA wt with erythrocytes labeled with the lipid marker R18 and the content marker calcein. 24 hours after transfection cells were treated with trypsin to cleave HA and incubated with double-labelled erythrocytes. Unbound erythrocytes were washed off and cells were treated for 5 min with pH 5, neutralized and monitored with a fluorescence microscope in the calcein channel and R18 channel. (A) Fluorescence microscopy (40x magnification) show diffusion of calcein (green) into the cytoplasm and of R18 (red) into the plasma membrane of cells expressing HA YKLW4A. Mock: untransfected cells do not show tight binding of erythrocytes to cells. (B) Comparison of the number of fusion events recorded for HA wt and HA YKLW4A at 20x magnification. Note that the number of erythrocytes is roughly the same in both microscopic fields, but cells expressing HA YKLW4A exhibit less hemifusion or full fusion. (C) Three examples of cells expressing HA YKLW4A that exhibit after acidification binding of erythrocytes that neither underwent hemifusion (diffusion of red dye into the plasma membrane) nor full fusion (diffusion of green dye into the cell’s interior, not displayed). (D) Relative quantification of individual fusion events for cells expressing HA wt or HA YKLW4A. In three transfections at least 100 cells with at least two bound erythrocytes were selected and (based on the result in the fluorescence channel) grouped into one of three categories, either unfused (=binding), hemifused or fully fused to erythrocytes. The total number of counted cells was normalized (=100%) and the percentage of events in each category was calculated and is displayed as the mean ±standard deviation.

Furthermore, cells expressing HA YKLW4A show one peculiarity not observed for HA wt, namely erythrocytes tightly bound to cells after acid treatment that did not pass through either hemifusion or pore formation, i.e. no diffusion of R18 into the plasma membrane (see Fig. 9C for three examples) and of calcein into the cell’s cytoplasm (not shown) did occur. We then selected cells with at least two bound erythrocytes and determined in the fluorescence channel whether the erythrocytes are unfused, hemifused or fully fused to the cells. We then calculated the percentage of individual fusion steps for HA wt or HA YKLW4A; the total number of selected cells was normalized to 100%. Cells transfected with HA wt did not show “binding” of erythrocytes after acidification, 58% of bound erythrocytes only passed the hemifusion step and 42% exhibit full fusion (Fig. 9D). Cells transfected with HA YKLW4A revealed a similar percentage of fully fused erythrocytes (45%), but reduced percentage of hemifused (28%) and 22% of bound, but unfused erythrocytes. Thus, we conclude that HA YKLW4A exhibits mainly a defect in hemifusion and probably also in another step that precedes lipid mixing.

## DISCUSSION

### Cholesterol binding to HA

In this study we show for the first time that HA interacts with cholesterol. We demonstrate that complete exchange of the cholesterol consensus motif YK…LW in a group 2 HA (H7 subtype) by alanine greatly reduces (>50%) photo-crosslinking of a cholesterol analog to HA (Fig. 2). This was demonstrated by labeling transfected cells with photocholesterol and thus for HA embedded in its natural lipid environment where photocholesterol must compete with other membrane lipids for the binding site in HA. Furthermore, a similar result was obtained for purified HA immunoprecipitated from cell lysates excluding the possibility that the stronger labeling of HA wt is due to its integration into cholesterol-enriched nanodomains. HA with an exchange of two consecutive amino acids YK and LW by alanine did not reveal reduced labeling with photocholesterol (Fig. 2) suggesting that the residues synergistically interact with cholesterol. However, note that photocrosslinking is a qualitative tool to measure the cholesterol affinity of a protein and may not represent equilibrium concentrations of protein-cholesterol complexes. To more precisely determine the amino acids in HA which contact cholesterol more sophisticated methods, such as NMR or crystallography are required (36).

Our data are at odds with a recent report showing by high-resolution secondary ion mass spectrometry that HA clusters at the plasma membrane of transfected cells were not enriched with cholesterol (59). However, the observation was made with an H2 subtype HA belonging to the phylogenetic group 1 that does not contain the YKLW motif. Although HA clusters at the plasma membrane have been observed both for group 1 (e.g. H2, (6)) and group 2 HAs (e.g. H3, H7, (7, 9, 27)) their mechanism of clustering might be different. Interestingly, protease digestion experiments and molecular dynamics simulations revealed that the TMR of group 2 HAs have a more compact and protease-resistant quaternary structure compared to the TMR of group 1 HAs (39).

The CCM is not only present in the consensus sequence of HAs of subtypes H3, H4, H7, H10, H14 and H15 (Fig. 1B), the amino acids YK…LW are also completely (99%) conserved within each subtype (60, supplementary file 1) arguing in favor for an essential role. Indeed, infectious virus particles could not be rescued if the CCM is completely exchanged; viral titers are lower if two amino acids are replaced. The mutant viruses are rapidly outgrown by wild-type indicating that they have a comparative fitness disadvantage (Fig. 3). It is thus safe to conclude that the CCM (and thus its interaction with cholesterol) is essential for Influenza virus replication, affecting both virus assembly and its cell entry via membrane fusion as discussed next.

Group 1 HAs do not possess the YK…LW motif, but contain the fairly conserved motif Y-K/Q…I-Y which also corresponds to the CCM defined for 7TMR (F/Y-R/K-I/V/L-Y/W). In addition, various rather loosely defined cholesterol recognition motifs exist that share a similar pattern of basic, aromatic and large hydrophobic amino acid (34). However, mutations at the boundary between the linker region and external part of the TMR do not retard transport of group 1 HAs to the plasma membrane (51). Note also that a tyrosine which is important for the recently determined structure of the TMR region of a group 1 HA (residue 18 in Fig 1a, (42)) is not conserved in group 2 HAs suggesting that both phylogenetic HA groups might exhibit different structures in this region.

### The role of the CCM for apical transport of HA and for virus assembly and budding

Mutations in the CCM did not have an effect on apical budding of virus particles from polarized cells and did not affect transport of HA to the apical membrane (Fig. 4). Although signals for apical transport of HA are located in the TMR they do not overlap with signals that mediate inclusion of HA into detergent-resistant membranes a surrogate marker for association with cholesterol enriched nanodomains (52).

However, the cholesterol content of mutant virus particles is significantly decreased by 10% (mutant LW2A) to 20% (YK2A, see Fig. 6). Assuming that the membrane of one spherical Influenza virus particle contains a total of 300.000 lipid molecules (as calculated for HIV particles that have the same size and hence lipid surface area (61) and ∼ 50% (=150.000) are cholesterol (5), a decrease of 10-20% is equivalent to 15.000-30.000 cholesterol molecules. An average Influenza virus particle contains 300-500 trimeric HA spikes (as determined by Cryo-EM, (62, 63)) and thus at most ∼ 1500 cholesterol binding sites are available. Based on this estimation it is evident that the 10-20% reduction in the cholesterol content cannot be explained by a stoichiometric (1:1) binding of cholesterol to HA. Instead, a cooperative effect must be involved; the mutation in the CCM decreases the cholesterol content by 10-20 cholesterol molecules per trimeric HA spike. One cholesterol molecule in direct contact with HA’s CCM recruits (possibly by lipid-lipid interactions) other steroid molecules into virus particles. This assumption is supported by crystal structures of the β-adrenergic and other 7TM-receptors that exhibit two (or more) parallel aligned cholesterol molecules, but only one interacts directly with the CCM motif (35).

10-20 cholesterol molecules would be sufficient to encase the outer part of transmembrane domain of one trimeric HA spike if we assume an α-helix and cholesterol diameter of 1 and 0.5 nm, respectively. Such a lipid shell has been postulated to target transmembrane proteins to lipid domains or induce the formation of domains in a membrane that is poised to do so but is not yet phase-separated (64, 65). Accordingly, the HA mutants YK2A and LW2A revealed reduced fluorescence resonance energy transfer (FRET) with a double-acylated raft-marker in transfected cells (37) and exchange of three amino acids at the beginning of the transmembrane region prevents raft-dependent clustering of H3-subtype HA at the plasma membrane (26). Mistargeting of HA might cause budding of virus particles through the (cholesterol-depleted) bulk phase of the plasma membrane. In accordance, the reduction of 10-20% cholesterol molecules in mutant virus particles is equivalent to a total cholesterol content of 45%, which corresponds to the cholesterol content determined for the whole apical membrane of polarized MDCK II cells (5).

Mutations in the CCM do not only affect the lipids in the viral membrane, but also reduce incorporation of HA (relative to M1 and NP) into particles. This was evident not only with viruses grown in MDCK II cells, but also (but less distinct) in embryonated eggs. The effect was most pronounced if the two amino acids LW in the TMR were exchanged (Fig. 5). A similar observation was made for H3 subtype HA from the filamentous Udorn strain having an exchange of the amino acids WIL at the beginning of the TMR (26). A priori one would rather assume that the HA content of mutant virus particles increases if viruses bud through the bulk phase of the plasma membrane but all other viral proteins are still mainly targeted to the original virus assembly site. However, assembly of viral proteins at the plasma membrane is a complicated process dependent on intrinsic signals in viral proteins as well as (partly transient) protein-protein interactions (8, 11, 12).

The mutation LW2A had the strongest effect on replication of virus in cell culture (Fig. 3B, 4A) and on HA incorporation into virus particles (Fig. 5), whereas the cholesterol content was reduced in the mutant YK2A to a larger extent compared to LW2A (Fig. 6). Note, however, that the data on the composition of virus particles show considerable variation between experiments, which is at least partly due to the pleomorphic nature of Influenza viruses. A recent study showed that even genetically homogenous virus particles released from a single infected cell show enormous variation in size and protein composition, i. e. the copy number of individual proteins vary up to 100 fold between virions (66). This low-fidelity assembly process makes it complicated to more precisely determine the effect of mutations on the morphology of Influenza virus particles.

An open question is the functional relationship between the two intrinsic nanodomain targeting signals in HA, S-acylation at cytoplasmic cysteine residues and the cholesterol-binding amino acids at the beginning of the TMR. Removal of only one signal is sufficient to perturb raft association of HA (27, 29, 30). Both signals are essential for virus replication; their complete removal prevented creation of recombinant virus particles (this study and (18, 60, 67)). Viruses with partially deleted signals could be generated but revealed lower titers and defects in virus budding and membrane fusion (18, 26, 67). Otherwise, effects of mutating the two nanodomain-targeting signals are different. Removal of the acylation sites does not retard intracellular transport of HA (31, 68) and does not change the lipid composition (cholesterol content) of the viral membrane, at least not if virus-like particles were analyzed (69). How the functions of both raft-targeting signals interact to define the viral budding site remains unclear.

However, the interaction of HA with cholesterol does not necessarily occur only at the plasma membrane since the HA mutants exhibit strongly retarded transport through the Golgi (37), the part of the exocytic pathway in which the cholesterol concentration successively increases from ∼5% (ER) to ∼40% (plasma membrane) (58).

### The role of the CCM for membrane fusion

With three different assays using either virus particles or HA-expressing cells we show that mutations in the CCM of HA decrease both the kinetics and the extent of HA’s fusion activity. A similar result was reported for H3-subtype HA that contains mutations at three amino acids at the beginning of the transmembrane region (26).

One contributing factor might be the reduced amount of HA at the surface of transfected cells (37) and in virus preparations (Fig. 4). However, we (at least partly) corrected for the latter by adjusting wild type and mutant viruses to the same HA-titer prior to the fusion assay. Thus, it is likely that the fusion defect is not (only) due to lower numbers of HA molecules at the fusion site, but a direct consequence of the mutations in the CCM of HA.

The R18 lipid mixing assay with mutant virus particles YK2A and LW2A as well as quantification of individual fusion events of HA YKLW4A-expressing cells revealed that the mutations in the CCM mainly affect the stage of lipid mixing (Fig. 7-9). Cells expressing HA YKLW4A showed another peculiarity, namely unfused erythrocytes still bound to their surface after low pH treatment (Fig. 9). In principle, this could be a fraction of HA YKLW4A molecules not activated by the low pH treatment and hence still bound in its pH 7 conformation to sialic-acid containing receptors on red blood cells. Alternatively, this HA YKLW4A fraction might have executed the first conformational change but has not completed the refolding step that causes fusion between viral and cellular membranes (1). In that case the fusion peptide has been exposed and inserted into the membrane thereby stabilizing the interaction between HA-expressing cells and erythrocytes at acidic pH. Our observation of tightly bound, but unfused erythrocytes to HA-expressing cells might correspond to early stages of HA-mediated membrane fusion recently observed by Cryo EM, e.g. HA-bridging, membrane dimpling and/or tightly docked membrane interfaces (70, 71). In any case, it suggests, that HA YKLW4A also has a defect in a membrane fusion step prior to lipid mixing.

The reduced fusion activity of HA might be due to a global effect of the mutations in the CCM on the viral membrane. Virus particles exhibit reduced cholesterol content, and this might profoundly affect biophysical properties of the membrane beneficial for fusion (22, 23, 25, 56)). Indeed, when we loaded mutant virus particles with additional cholesterol the hemolysis activity increased (Fig. 7D). Nevertheless, cholesterol-loaded mutant virus particles revealed hemolysis values well below those determined for FPV* wt without additional cholesterol loading although their cholesterol content is now roughly the same (Fig. 7C). Note also that the HA mutant YK2A has the lowest cholesterol content in the viral membrane (Fig. 6B), but HA LW2A exhibits the largest effect on virus infectivity and its hemolytic activity (Fig. 3+7). Thus, the fusion defect is most likely not due to a general disadvantageous property of the viral membrane, such as disturbed liquid phase separation (raft-formation) and/or membrane ordering. We therefore rather prefer a model where a local interaction between cholesterol and the TMR of HA affects membrane fusion as proposed recently (57). One might envision that the CCM recruits cholesterol from the inner to the outer leaflet of the viral membrane, especially prior to hemifusion. The shape of cholesterol (small headgroup, large hydrophobic tail) is beneficial for the formation of a highly bended membrane intermediate, but only if cholesterol is located in the external leaflet, and not in the internal leaflet of the viral membrane (21, 72). Alternatively, cholesterol binding to the CCM (or more generally the amino acid exchanges we introduced) might affect the flexibility of HA’s linker region which may be important to facilitate the pH-dependent changes in HA conformation required for membrane fusion as recently suggested (42).

In sum, mutations in the cholesterol consensus motif of a group 2 HA affect various functionalities of the protein; its transport along the exocytic pathway, raft association at the plasma membrane (37), incorporation of cholesterol and HA into budding virus particles and virus entry via membrane fusion, especially lipid mixing and probably a preceding step. High resolution Cryo-EM of full length of a group 2 HA embedded in a membrane (similar to the one published for a group 1 HA (42)) might be helpful to determine the structure of the cholesterol binding pocket as one prerequisite to develop a small molecule that inactivates the virus.

## ACKNOWLEDGEMENTS

This work was supported by the German Research Foundation (SFB 740 TP C3) and by the Human Frontiers Science Program. Bodan Hu is recipient of a PhD fellowship from the China Scholarship Council (CSC). The funders had no role in study design, data collection and interpretation, or the decision to submit the work for publication. We thank Ralf Wagner and Hans-Dieter Klenk (Virology, Marburg) for providing the reverse genetics system used and Kai Ludwig (BioSupraMol, Chemistry and Biochemistry, FU Berlin) for performing electron microcopy.

