## supplementary file 1 for "Cholesterol binding to the transmembrane region of a group 2 HA of Influenza virus is essential for virus replication affecting both virus assembly and HA’s fusion activity"

**Synthesis of click-photocholesterol**

**Outline of the synthesis:** The synthesis starts with 3-TBDMS-26-hydroxycholesterol (i.e. compound 14 of (2)). After protection of the 26-hydroxy group with TBDMS-Cl, the double bond was converted into the 6-keto group by treatment with $BH_3$ followed by hydrolysis of the borane and oxidation with PCC. The protecting groups were removed and the resulting 3,26-dihydroxy-6-keto cholesterol was converted into the 6,6'-diazirine using the classical $NH_3/H_2N\text{-}OSO_3H/I_2$ method. By a sequence of protection-deprotection reactions, this diazirine was converted into the 3-TBDMS-derivative with a free 26-OH group. This primary alcohol was oxidized to the aldehyde and converted to the alkyne by treatment with Bestmann-Ohira reagent. Finally, the TBDMS-group at the 3-hydroxy group was removed to give the final 6,6'-Azi-25-ethinylcholesterol.

3,26-bis-(TBDMS-oxy)-cholesterol (**1**): 750 mg (1.5 mmol) 3-TBDMS-oxy-26-hydroxycholesterol were stirred in 10 ml THF with 3 eq. of TBDMSCl and 6 eq. of imidazole for 4 h at 30°C. THF was evaporated and the residue purified by silica column chromatography in hexanes/ethyl acetate 6/1 to yield 840 mg product.

3,26-bis-(TBDMS-oxy)-6-keto-cholesterol (**2**): 840 mg **1** were dissolved in 2 ml THF. 5 ml 1 M $BH_3$ in THF were added and the solution left at room temperature for 70h. Under stirring, 1 ml water, 20 ml ether and 5 ml brine were added subsequently. The lower phase was removed and the upper phased dried with sodium sulfate. The solvent was removed and to the residue were added 10 ml dichloromethane, 1 g powdered molecular sieve and 1.5 g pyridinium chlorochromate. The mixture was refluxed with occasional TLC control of the progress of oxidation. After 2h the solvent was evaporated and the residue extracted several times with hexanes/ethyl acetate 6/1. The extracts were combined and evaporated and the residue purified by silica column chromatography in hexanes/ethyl acetate 6/1 to yield 620 mg product.

3,26-dihydroxy-6-keto-cholesterol (**3**): 620 mg of **2** were dissolved in 10 ml THF followed by addition of 1 ml 6 N HCl and stirring for 1 h at 45°C. The solvent was removed at reduced pressure and the residue purified by silica column chromatography in hexanes/ethyl acetate/methanol 10/10/1 to yield 350 mg product. 1H-NMR in $CDCl_3$/MeOH shows the absence of the double bond proton at C-6 and the shift of the C-19 methyl protons to 1.05 ppm, see spectrum 1.

3,26-dihydroxy-6,6'-azi-cholesterol (**4**): 320 mg of **3** (0.75 mmol) were dissolved in 40 ml methanol. The solution was cooled with ice and ammonia was bubbled into this solution until saturation was reached. A solution of 2 mmol hydroxylamine-O-sulfonic acid in 2 ml methanol was added followed by three hours stirring while the temperature was allowed to increase to room temperature. Precipitated salts were filtered off and the residue was concentrated to 10 ml by evaporation at reduced pressure. Then 20 ml methanol and 1 ml trimethylamine were added followed by addition of solid iodine under stirring until the iodine color persisted (ca. 120 mg iodine). The solvent was evaporated at reduced pressure and the residue purified by silica column chromatography in hexanes/ethyl acetate/ethanol 50/50/2 to yield 80 mg product. 1H-NMR in $CDCl_3$/MeOH shows the the shift of the C-19 methyl protons to 1.05 ppm, see spectrum 2.

3-TBDMS-oxy-26-hydroxy-6,6'-azi-cholesterol (**5**): 80 mg of **4** were dissolved in 1 ml THF. After addition of 100 µl 2,6-lutidin the solution was cooled to 0°C and a 2M solution of acetylchloride in dichloromethane was added in portions of 50 µl until about 90% of the educt were consumed as indicated by TLC in hexanes/ethyl acetate/ethanol 50/20/1. By TLC, the mixture consisted of 10% educt, 70% 3-OH-26-acetyl protected, 5% 3-acetyl-26-OH- and 15% 3,26-bis-acetyl- protected material. The reaction was quenched by addition of 50 µl methanol followed by 10 ml of ether, 1 ml of water and 1 ml saturated NaHCO3. After shaking, the ether phase was collected, dried and evaporated. The residue was dissolved in 1 ml THF and 80 mg imidazole and 100 mg TBDMSCl wer added. The mixture was stirred overnight. TLC control showed nearly complete conversion to three bands at a ratio of 20/70/10 representing bis-TBDMS, 3-TBDMS-26-acetyl and bis-acetyl-protected material. The solvent was removed and the residual mixture treated with 1 M KOH in 2 ml MeOH at 45°C for 1h to remove the acetyl group on O-26. Now, TLC control in hexanes/ethyl acetate/ethanol 50/20/1 showed three bands corresponding to bis-TBDMS, 3-TBDMS-26-OH and 3,26-di-OH compounds. The mixture was separated by silica column chromatography in a) hexanes/ethyl acetate 40/10, b) hexanes/ethyl acetate/ethanol 50/50/1, c) hexanes/ethyl acetate/ethanol 50/50/1 to obtain the 3-TBDMS-26-OH compound and the two side products. The bis-TBDMS-form was deprotected with HCL in THF, combined with the dihydroxy-form and recycled for a second round of protecting group chemistry as described, yielding a total of 65 mg 3-TBDMS-oxy-26-hydroxy-6,6'-azi-cholesterol.

25-ethinyl-6,6'-azi-cholesterol (**6**)  To 65 mg of **5** were added 200 mg powdered molecular sieve,  2 ml dichloromethane and 120 mg pyridinium chlorochromate. After 30 min, the solvent was removed at reduced pressure and the residue purified by silica column chromatography in hexanes/ethyl acetate 4/1 to yield 60 mg of the 26-aldehyde that was directly dissolved in 2 ml methanol/THF 3/1. 200 mg potassium carbonate and 180 mg Bestmann-Ohira reagent were added and the mixture was stirred at room temperature for 2h. The solvent was evaporated and the residue purified by silica column chromatography in hexanes/ethyl acetate 6/1 to yield 48 mg of the 3-protected alkyne. This intermediate was dissolved in 1 ml THF and 100 µl 25% HCl were added followed by stirring for 30 min at 40°C. The solvent was evapotated at reduced pressure and the residue purified by silica column chromatography in hexanes/ethyl acetate/ethanol 60/30/4 to yield 36 mg final product.

**Analysis:** a 1H-NMR spectrum in chloroform was recorded along with spectra of authentic cholesterol, 6-photocholesterol (3) and click-cholesterol (1). The spectra clearly show the characteristic changes expected for the 6,6'-diazirine ring in photocholesterol and the 25-ethinyl function in click-cholesterol. The click-photocholesterol shows a hybrid spectrum with both the characteristic elements of the diazirine and the ethinyl groups. The UV-spectrum demonstrates the presence of the diazirine ring by the characteristic doublet peak at 353/371 nm, e = 79 $mol^{-1}cm^{-1}$.

HR-MS1 of **6** in the presence of ammonium acetate: m/z = 442.3796 $[M + NH_4]^+$ (theor. 442.3792), 407.3425 $[M + H – H_2O]^+$ (theor. 407.3421) MS2 of 442.4: m/z = 425.3779 $[M + NH_4 – H_2O]^+$, 414.3732 $[M + NH_4 - N_2]^+$ (theor. 414.3730)

### Figures:

1H-NMR of 6,6'-Azi-26-hydroxycholesterol in $CDCl_3$/MeOH

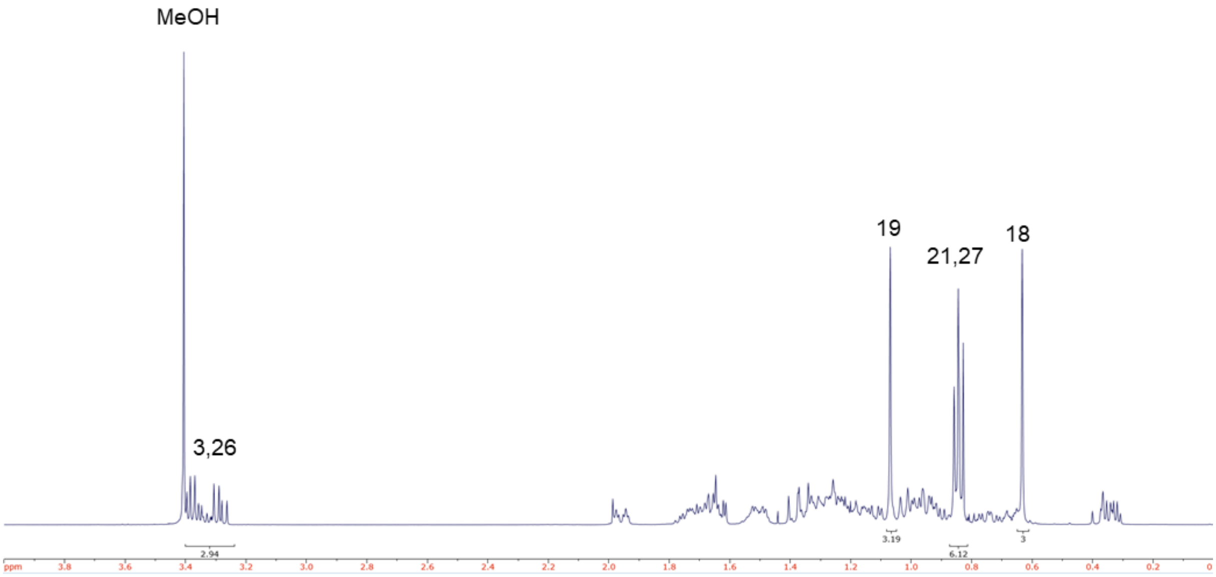


**Spectrum 1**

1H-NMR of 6,6'-Azi-25-ethinylcholesterol in $CDCl_3$

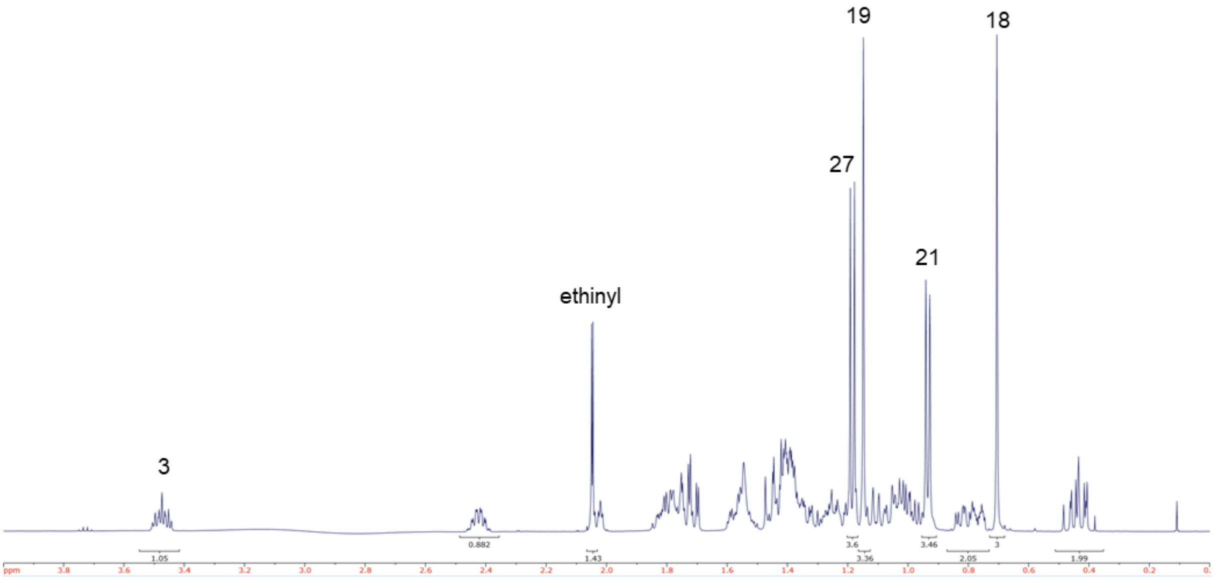


**Spectrum 2**

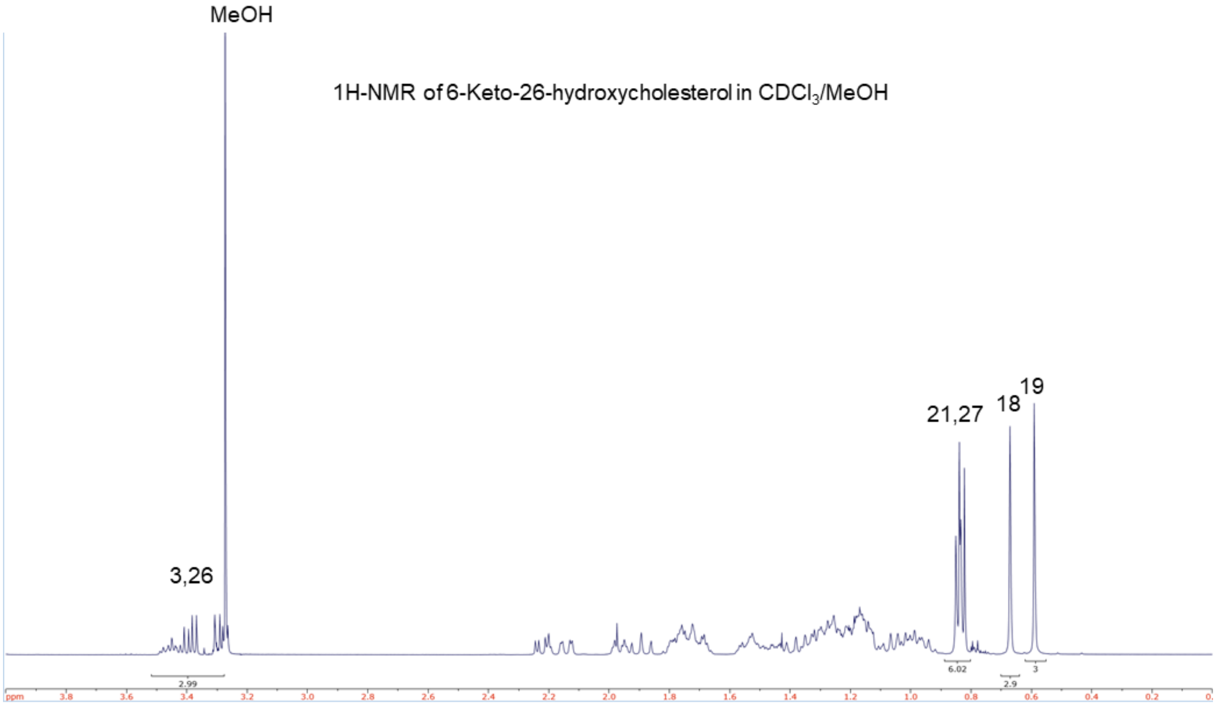


**1H-NMR spectrum of 6-Keto-26-hydroxycholesterol in chloroform methanol**

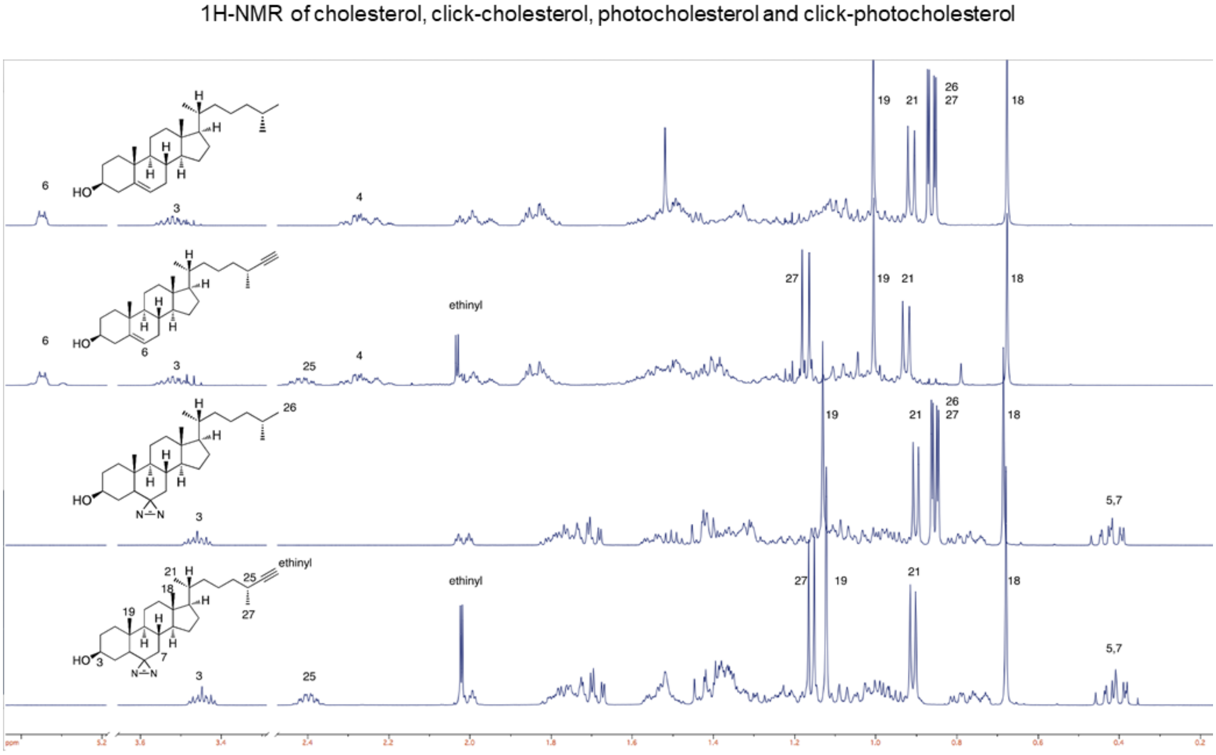


**1H-NMR spectrum of cholelesterol, click-cholesterol, photocholesterol and click-photocholesterol in chloroform**
